# Longitudinal wastewater sampling in buildings reveals temporal dynamics of metabolites

**DOI:** 10.1101/2019.12.16.870576

**Authors:** Ethan D. Evans, Chengzhen Dai, Siavash Isazadeh, Shinkyu Park, Carlo Ratti, Eric J. Alm

## Abstract

Direct sampling of building wastewater has the potential to enable “precision public health” observations and interventions. Temporal sampling offers additional dynamic information that can be used to increase the informational content of individual metabolic “features”, but few studies have focused on high-resolution sampling. Here, we sampled three spatially close buildings, revealing individual metabolomics features, retention time (rt) and mass-to-charge ratio (mz) pairs, that often possess similar stationary statistical properties, as expected from aggregate sampling. However, the temporal profiles of features—providing orthogonal information to physicochemical properties—illustrate that many possess significantly different *feature temporal dynamics* (fTDs) across buildings, with rapid and unpredictable deviations from the mean. Internal to a building, numerous and seemingly unrelated features, with mz and rt differences up to hundreds of Daltons and seconds, display highly correlated fTDs, suggesting non-obvious feature relationships. Data-driven building classification achieves high sensitivity and specificity, and extracts building-identifying features with unique dynamics. Analysis of fTDs from many short-duration samples allows for tailored community monitoring with applicability in public health studies.

---

Wastewater sampling presents a means to monitor the general health^1^, chemical exposure^2,3^, and size^4^ of a population in a rapid and noninvasive manner. Many studies have been performed at wastewater treatment plants (WWTPs), as these sites are relatively easy to sample, and yield aggregate information on large populations, e.g. from an entire city^2^. As an example of the correlations that can be captured by such studies, an increase in antipsychotics, antidepressants and other therapeutic drugs was observed in wastewater between 2010 and 2014 in Athens, Greece during a time of significant economic turmoil in the country5. Given the proper sampling and analysis methods, wastewater can provide meaningful, community-specific public health information.

Most wastewater epidemiology and metabolomics studies have focused on the aggregate chemical load on large populations, typically using targeted metabolomics to acquire highly sensitive, context-specific information about select small molecules present at WWTPs. City- and country-wide studies have focused on monitoring licit^5,6^ and illicit drugs^7–9^, including sports doping agents^10,11^, and have used these results to estimate public drug consumption^12^. Targeted applications have ranged from monitoring stress-related molecules13, plasticizers^3,14^, and pesticides^2^ to metabolites associated with alcohol^15^ and tobacco^16^; general population biomarkers^17^; and the environmental release of pharmaceuticals^18^. In addition to chemical identification, wastewater metabolomics can also be used to estimate population size^19,20^. However, aggregate analysis of large populations potentially misses public health-relevant information on temporal dynamics and sub-population characteristics.

There are multiple ways to incorporate temporal information in wastewater metabolomics that depend on sampling methods and location. One common route is to collect many composite samples, often over 24 hours, via extended continuous sampling^4,21,22^. This route is logistically easier, as it only requires a single site. However, the temporal component provides an averaged signal, even with multiple single day composite samples. An alternative is to perform close-to-source, periodic grab sampling or short continuous sampling without combination^23^. The second route often requires multiple locations and high sampling frequency (hourly to daily or near-daily), necessitating a large number of samples. Extensive sampling is required to alleviate the problem of signal noise and stochasticity associated with short sampling of small populations. However, a significant benefit of this approach is that it provides a precise temporal snapshot of the molecules present at a given time and, with longitudinal samples, temporal dynamics with minimal signal averaging. This sampling may provide information on individual contributions to community wastewater, and when compositional shifts or chemical exposure events occur.

We conducted a multi-month, untargeted metabolomics analysis of wastewater from three individual buildings to understand the information contained in high temporal resolution sampling of small populations. To study the temporal resolution needed to characterize a small population, we performed near-daily sampling for one month, followed by roughly biweekly sampling over two months from a multipurpose-use building (Building 1) and two residential buildings (Buildings 2 and 3). While through-time statistical values (mean intensities) of the features showed minimal differences between buildings, analysis of *feature temporal dynamics* (fTDs) uncovered extensive differences. Temporal feature clustering and modeling, internal to each building, revealed numerous groups of shared fTDs that often displayed random but large intensity fluctuations. These dynamics would likely be unobserved or averaged out using alternative sampling approaches. Similarity of fTDs suggested links from putative metabolites to unknowns as well as between features with drastically different mz and rt values, both within and across buildings. To extract additional building-distinguishing information, we trained interpretable machine learning (ML) models using daily feature profiles. Extending and generalizing our analysis methods, we found additional fTDs that correlated with those of select putative molecules, suggesting features for follow-up analysis.

## Results

### 1. Traditional statistical approaches do not capture the full temporal differences between buildings

Feature summary statistics (mean and standard deviation through time of feature ion intensities) provide a simple method to conceptualize and coarsely categorize feature stability. This stratification allows one to triage the features according to the research question of interest. Using this approach, we identified stable and unstable features that were generally similar between buildings. Further subcategorization and analysis of the unstable features revealed unique day-to-day dynamics, suggesting that summary statistics do not fully capture temporal dynamics that are essential components of a small population’s wastewater metabolome.

#### 1.1. Longitudinal multi-month sampling allows for temporal variation-based feature grouping

We observed two distinct groups of features during multi-month sampling of three spatially close buildings: temporally stable, and temporally unstable. Sampling occurred over three months, with the most dense sampling (multiple times per week) occurring in the first 3 weeks, followed by sampling approximately 1–2 times per week for the remainder of the period (Figure 1A). Liquid chromatography mass spectrometry (LCMS) produced 1440 features, only days for which LCMS data from all three buildings was available were used in subsequent analysis. The features were separated by through-time standard deviation; values < 3 were considered stable, and ≥ 3, unstable. The majority of stable features were stable in all three buildings (61% of features—Figure 1B). Similarly, though to a lesser extent, 40% unstable features were unstable across buildings, but larger fractions were uniquely unstable in one or two buildings (Figure 1B). A fraction of features were putatively identified at minimum reporting standard^24^ levels 2 or 3, while most were unidentified (level 4, see *Methods* for definitions).

**Figure 1.**
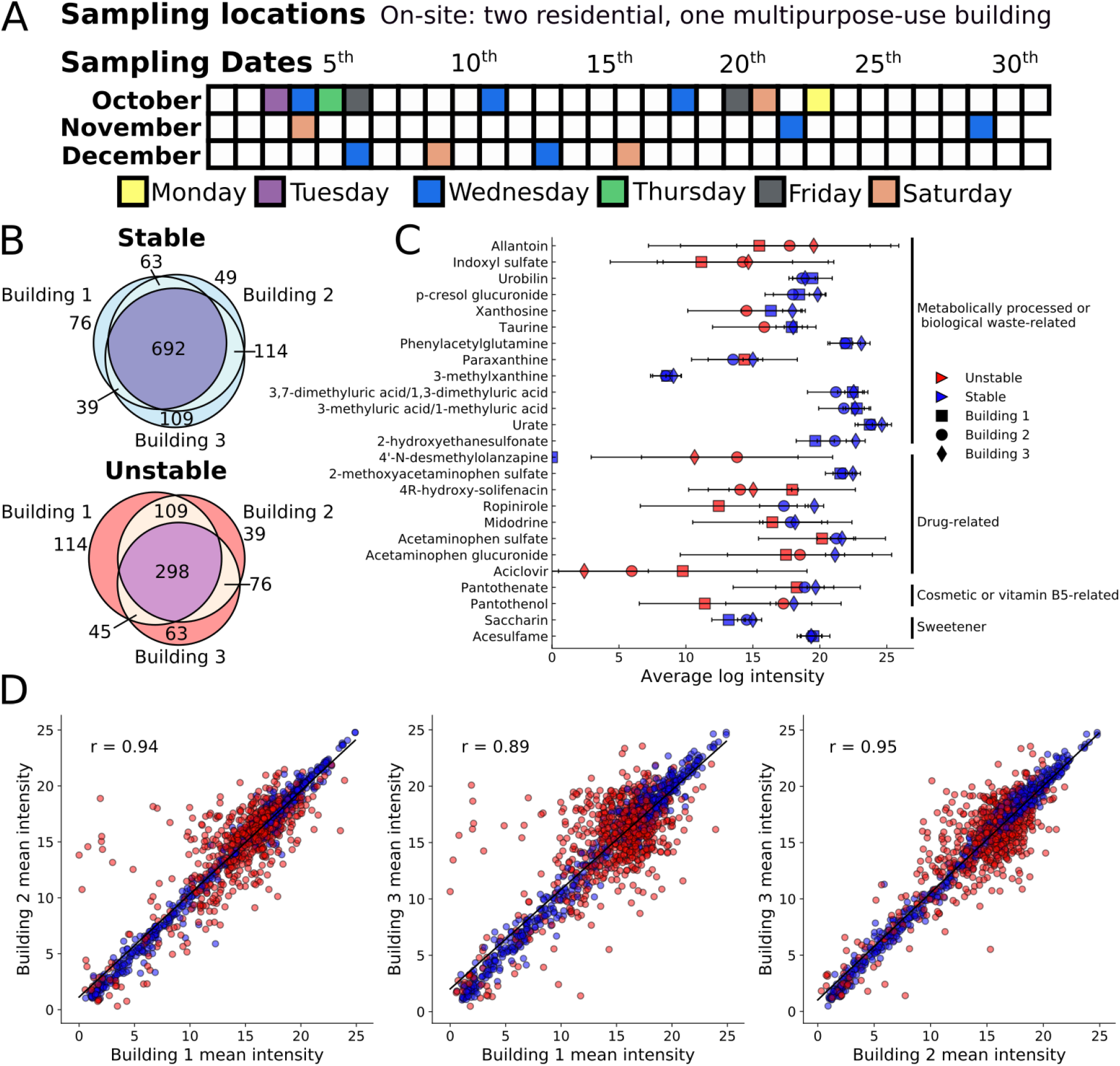
Multi-month sampling reveals numerous temporally stable and unstable features that generally possess comparable mean intensities across buildings. (A) Sampling timeline across three months for all buildings with sampling days filled with color. (B) Breakdown of the stable (through-time standard deviations < 3) and unstable metabolites and their overlap across the three buildings. (C) Average through-time mean log intensities and standard deviations of select features that were mapped to putative metabolites (level 2 identification); panel displays a selection of metabolites related to human activity, drugs, vitamin B5, and sweeteners. (D) Average through-time mean log intensity comparisons between all combinations of the three buildings. Red dots represent unstable features; blue dots represent stable features in at least one building.

Untargeted metabolomics revealed numerous putative human and human-associated small molecules, both in the stable and unstable categories, that displayed a range of through-time mean intensities, standard deviations, and building-to-building variability. We observed several chemicals and common metabolic products of human activity, including metabolites from caffeine metabolism (xanthine-based metabolites^25^), dietary tryptophan processing (indoxyl sulfate^26^) as well as expected urinary metabolites (urate^27^, phenylacetylglutamine^28^, and 2-hydroxyethanesulfonate^29^—Figure 1C). The majority of these metabolites were stable in all three buildings, with the exception of indoxyl sulfate, which was unstable across all three buildings. Putative drug-related metabolites displayed increased feature instability and primarily consisted of acetaminophen metabolites plus possible drugs for restless leg syndrome (ropinirole) and low blood pressure (midodrine). Putative artificial sweeteners (acesulfame and saccharin) appeared stable across the three buildings, but chemicals naturally found in humans as well as in many health and cosmetic products (pantothenate (vitamin B5) and its precursor pantothenol—Figure 1C) displayed high variability. A large number of the drug- and cosmetic-related features appeared unstable in many buildings, particularly in Building 1.

Through-time statistical analysis suggested that individual features often appeared at similar mean intensities for all three building-to-building comparisons—especially the stable features. Linear relationships were observed for feature intensity comparisons between all buildings (R=0.94, 0.89 and 0.95 for the Building 1-to-2, 1-to-3 and 2-to-3 comparisons, respectively—Figure 1D). While the high correlations were calculated using all features, the unstable features were more dispersed, and minimally correlated, between buildings. Assuming consistent ionization across samples, this suggests that many features exist at comparable average concentrations in buildings with different populations.

#### 1.2. Few temporally stable features are statistically different between buildings

Only 16 stable features were present at significantly different mean levels between the three buildings (false discovery rate, FDR, corrected Kruskal-Wallis, KW, P-value < 0.00001). Relative to Building 1, the majority of features appeared at lower mean intensities in Building 3, while those in Building 2 appeared at higher levels (Figure 2A). Of the 16 significant features, 13 were observed at higher levels in Buildings 2 and 3 relative to Building 1, in contrast to the general feature trends (Figure 2B). Unlike manually searching for specific metabolite types, statistical analysis found the urine metabolite 2-hydroxyethanesulfonate and sweetener saccharin among the handful of significant features (Figure 2C). Few commonalities were observed between these features as they appeared across a wide range of mz, rt, and intensity values from less than five to near 25.

**Figure 2.**
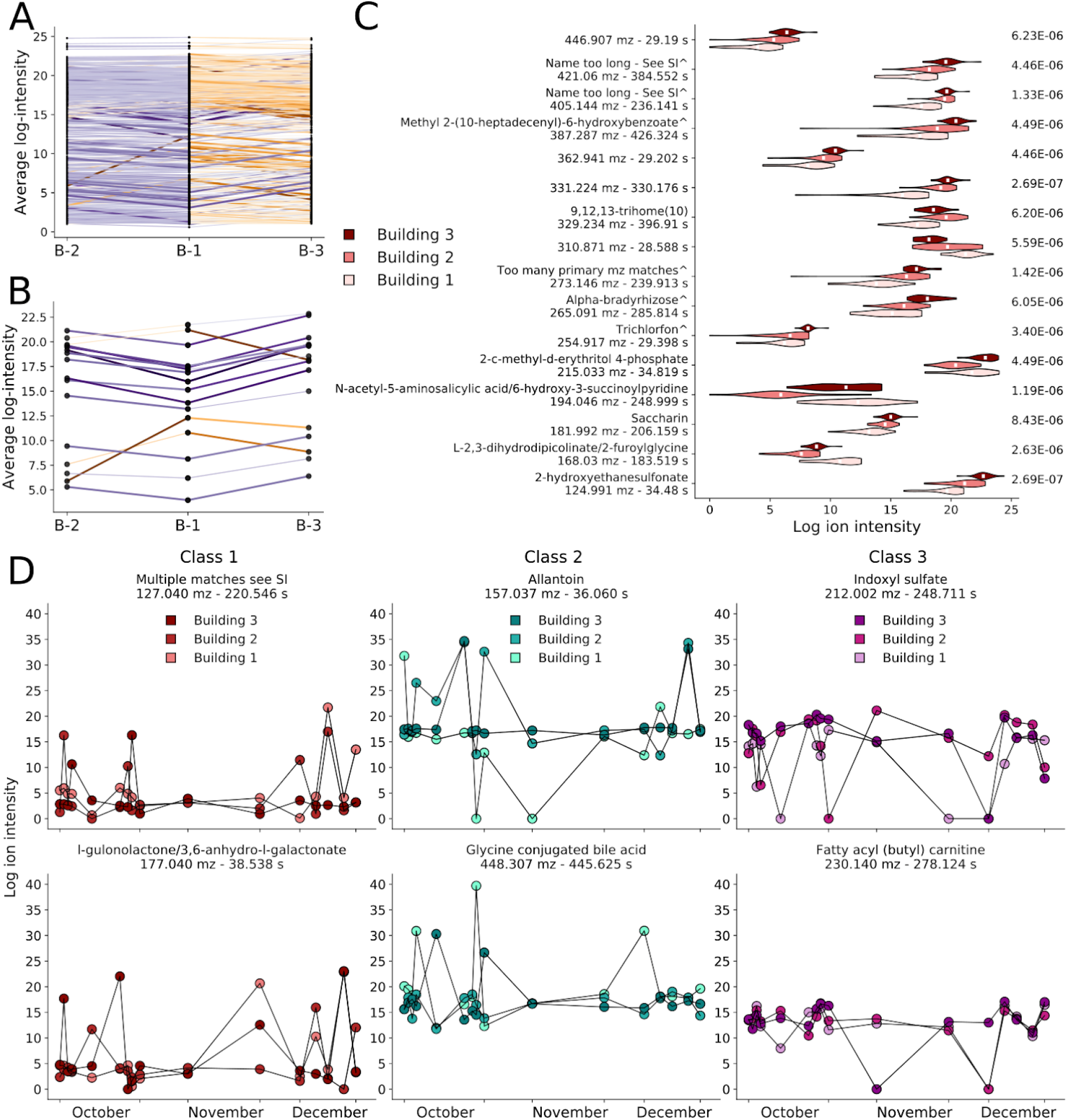
Only 16 out of 1142 stable features are statistically different between buildings, and unstable features display three temporal classes. (A) Average log intensity comparison of all stable features across buildings. Purple (orange) lines depict features at a lower (higher) intensity in Building 1 relative to the other building, larger absolute intensity differences correspond to darker lines. (B) Comparisons for stable features that are also statistically significant between buildings (FDR-corrected KW test, Q < 0.00001). (C) Violin plot of all 16 significant features depicting building-to-building differences and distributions. Names are levels 2 and 3 putative identifications (names ending with an ^ are level 3; see Table S2 for full names). Right column, associated Q-values. (D) Unstable features show three classes: class 1 features (left column) possess median log intensities < 5; class 2 features (middle column) show intensity spikes greater than the largest value recorded for all stable features (~26); and class 3 features (right column) were the remaining features that generally displayed mid-level intensities with occasional, rapid decreases (see Table S3 for names).

#### 1.3. Unstable features can be further split into three dynamics-based classes

Temporal analysis of the unstable features suggested three general temporal profiles, providing a more fine-grained classification, and a means for conceptualizing their dynamics (Figure 2D, Table S3). Class 1 contained features with a through-time median intensity less than five, with random spikes to higher intensity values. This class included metabolites that are typically observed at low levels in wastewater, but that are occasionally present at high levels. Class 2 unstable metabolites possessed high through-time intensity levels, and at least one, often unpredictable, spike that surpassed the largest ion intensity in all the stable features (≳ 26). Class 2 not only possessed allantoin, but also putative glycine conjugated cholic acids (Figure 2D, middle panels). Class 3 was comprised of features possessing mid-level log intensities with occasional, unpredictable dips in intensity to a (near) zero value. Representative examples of this class included indoxyl sulfate and fatty acyl (butyl) carnitines (Figure 2D, right panels). Temporal analysis departs from using solely through-time statistical parameters to describe a time-series; using only three simple groups of feature temporal dynamics (fTDs) highlighted significantly different profiles—this prompted a more comprehensive dynamics analysis.

### 2. Characterizing and modeling the dynamics of individual buildings with clustered fTDs

Temporal analysis supplied a more nuanced view of the features, and thus buildings, than binary stability types. Unsupervised clustering revealed a large diversity of dynamics in a community’s wastewater metabolome. Two modeling approaches illuminated the importance of using many short-duration samples to capture rapid fluctuations and a community’s chemical output at a given time. Fitting clusters with a Gaussian process (GP) suggested that time points supplied little information on each other. In addition, theoretical waste mixing showed substantial decreases in feature intensity dynamics when only a small number of additional waste streams were combined.

Differences between buildings were further highlighted by focusing on the temporal profiles of putative classes of molecules related to metabolism and lifestyle.

#### 2.1. Clustering fTDs uncovers groups of features with highly similar temporal dynamics

K-means clustering of individual building features displayed several prominent families that differed between buildings. One hundred clusters were used to group z-normalized features, providing highly similar intracluster dynamics, mostly within the range of −1 to 1 (Figure 3A). The high temporal sampling revealed that many clusters exhibited sudden, single-day spikes or drops in intensity (Figure 3A, dark blue and red regions). Additionally, many days displayed cross-cluster intensity correlations; for example in Building 1, on October 6^th^ the top 8 clusters showed similar intensities, and on Saturday, December 16^th^ the majority of clusters demonstrated a general drop in intensity for many features. When comparing across the three buildings, no obvious trends were observed in terms of mean cluster dynamics, which varied between buildings. The majority of clusters in all buildings were composed of features spanning a large range of mz and rt values (Figure 3A purple and green columns). Additionally, a relatively small number of temporal profiles (~50 clusters) accounted for 74–81% of the buildings’ features (Figure S1).

**Figure 3.**
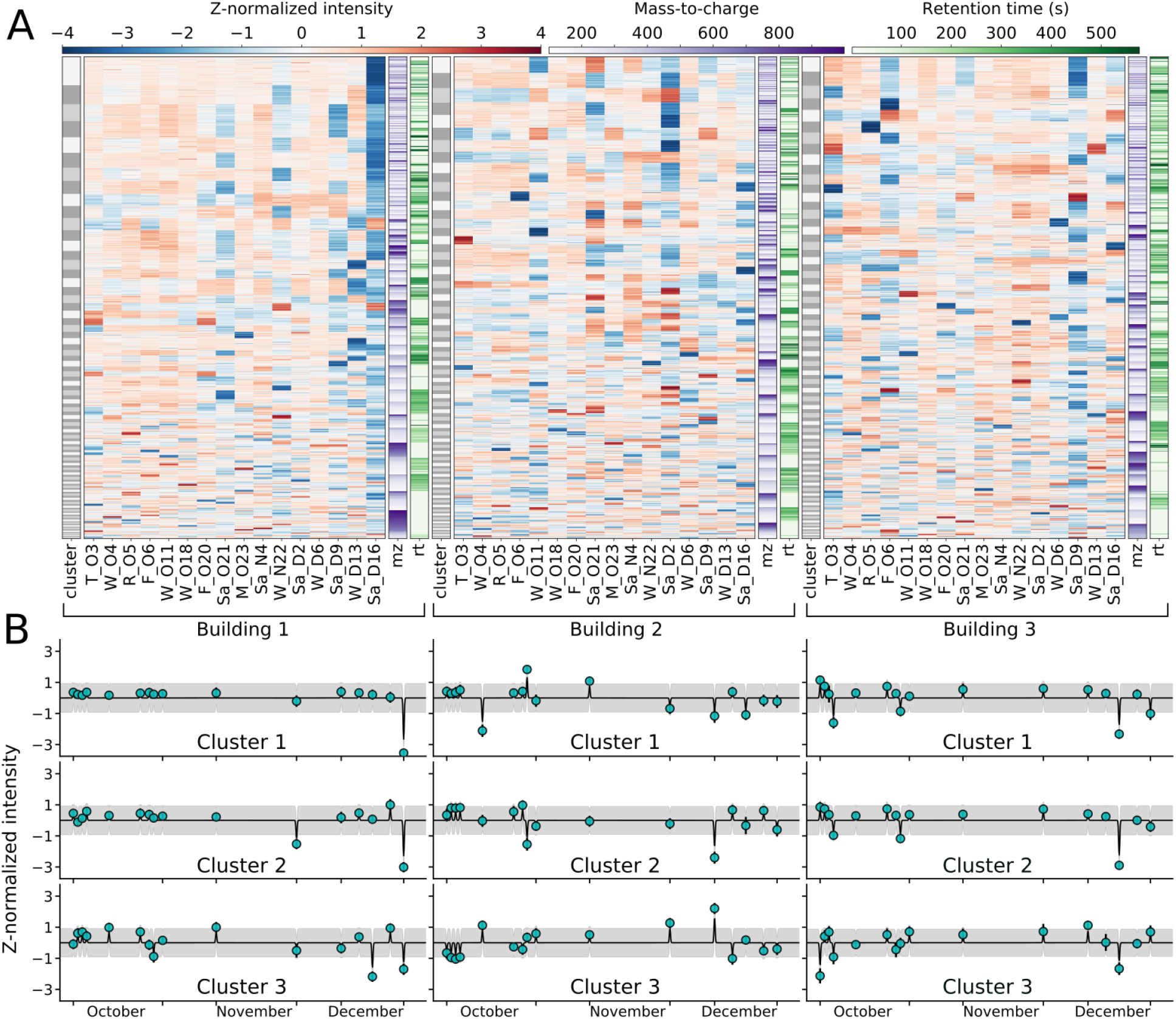
Individual buildings possess unique fTDs with most dynamics governed by a few clusters and rapid deviations from the mean. (A) Summary of all building data sets using only shared sampling days. Gray columns show the 100 clusters, with individual box heights corresponding to cluster size. The 16 red and blue columns depict single-day, z-normalized log intensities for each feature (individual rows) for the three buildings; purple and green columns show feature mz and rt respectively. Color keys shown above the plots. (B) Average cluster values (teal points) for the three largest clusters for each building, with each cluster fit to a GP. Model standard deviation in gray.

GP cluster modeling displayed minimal day-to-day predictive power, and rapid reversion following large deviations from the mean. As a previous 24-hour metabolomics analysis^23^ demonstrated strong diurnal patterns in wastewater, we used a small length scale parameter for the GP. Using a conservative 3-hour length scale, information decayed rapidly, providing no day-to-day predictive power. With this length scale, the GP models showed a rapid mean-reverting tendency following large perturbations, accompanied by increased predictive uncertainty (Figure 3B). GP modeling demonstrated the importance of high-frequency sampling and indicated that the biweekly sampling likely missed significant wastewater dynamics.

#### 2.2. Theoretical waste stream mixing suggests dynamics are lost with a small number of additional sources

The rapid and unique fluctuations observed in single-building, 3-hour samples were lost under simulations of slightly longer sampling periods, or upon combination with relatively few additional wastewater sources. We combined and clustered the data from the three buildings with a desired number of modeled waste streams, with each cluster fit by a GP (see *Methods*). Repeating this protocol with different numbers of combined waste streams suggested that major cluster dynamics are dampened when as few as tens of waste streams are combined, and lost with more than 50 (Figure 4). GP modeling of the combined data suggested that the GP uncertainty rapidly drops as additional waste streams are mixed. Under our modeling assumptions, features will be found at their expected values with high probability and minimal day-to-day variation (Figure 4A–E). To measure how the differences between clusters decrease with additional waste streams, we calculated the summed all-to-all cluster mean Euclidean distance. This demonstrated that cluster differences are effectively lost when waste from 50–100 sources is merged (Figure 4F). Molecules that are rarely observed or that come from a small number of sources may still display significant dynamics; however, the majority of features will generally be observed at statistical averages. This underscores the importance of short, upstream collection to obtain relevant public health information.

**Figure 4.**
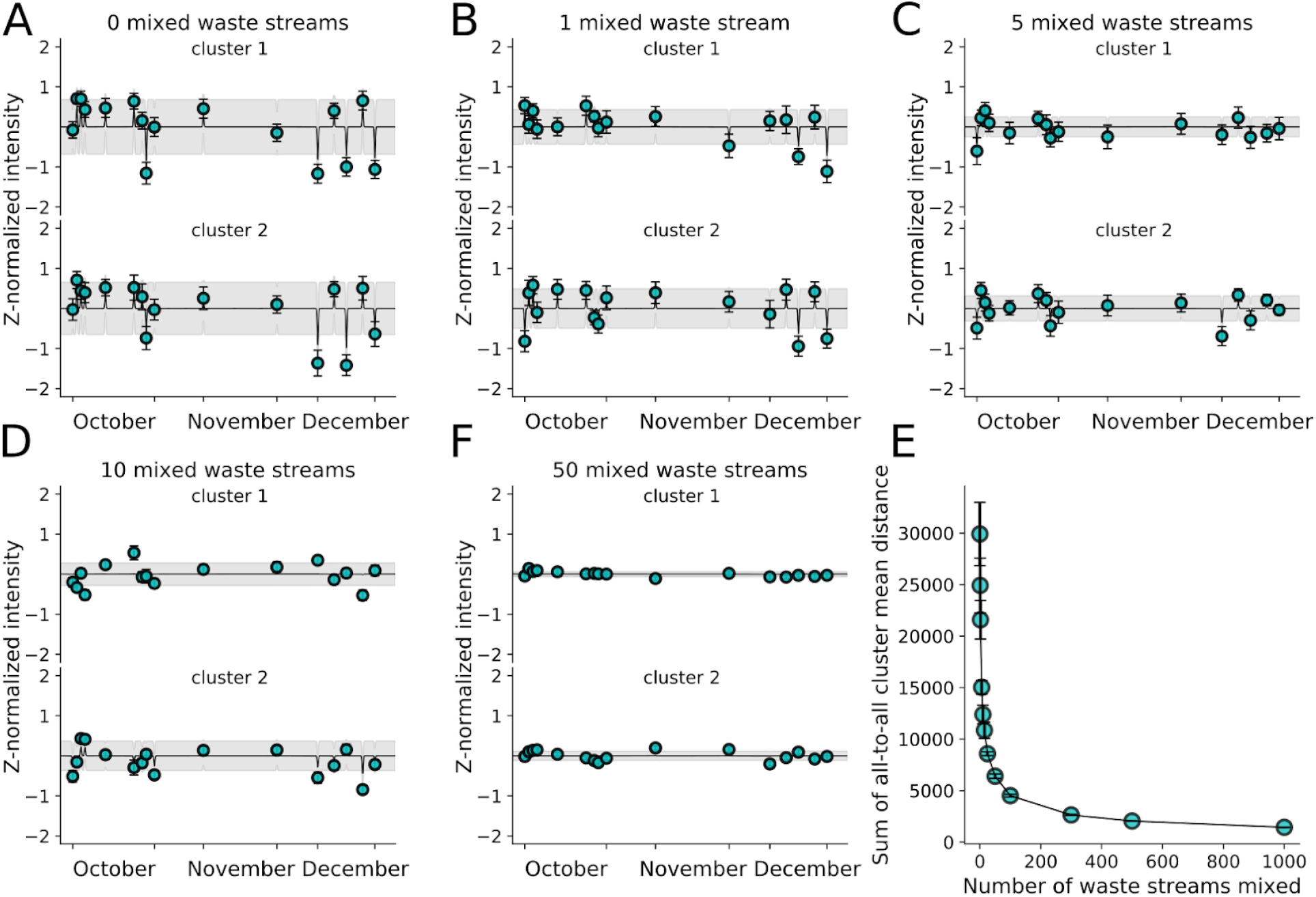
Large feature deviations from the mean are lost with the addition of relatively few waste streams. (A–E) Mixing of building waste with 0–50 additional simulated waste streams, with clustering and GP fitting (see *Methods*). The two largest clusters are plotted, with their standard deviations in gray. (F) Plot of the sum of Euclidean distances between all pairs of cluster centers for a given number of mixed buildings. Error bars represent the standard deviation following 10 repeats.

#### 2.3. Frequent sampling allows for tracking dynamics of human-related metabolite groups

The three buildings displayed different dynamics for specific human metabolic and health-related putative metabolite classes. Metabolite groups studied included glucuronide-modified compounds^30^, caffeine-related metabolites^25^, biologically modified acetaminophen^31^, along with glucoside conjugated molecules (Figure 5, Figure S2, and Table S4 for full names). These human-associated putative groups, possessed diverse temporal dynamics in each group and building. For instance, many of the features showed large changes across buildings; however, the days on which the specific dynamics occurred differed for each given putative metabolite. Within each building, select putative metabolites from each group displayed similar temporal dynamics, perhaps due to similar biochemical processing (e.g. different metabolites of acetaminophen). However, not all temporal profiles in a group were always similar, for instance the October levels of many glucuronides and caffeine metabolites displayed pronounced differences. While the ability to identify additional features is required for larger, targeted chemical tracking, this analysis highlighted the potential of high-frequency wastewater sampling to monitoring groups of health and lifestyle-related compounds.

**Figure 5.**
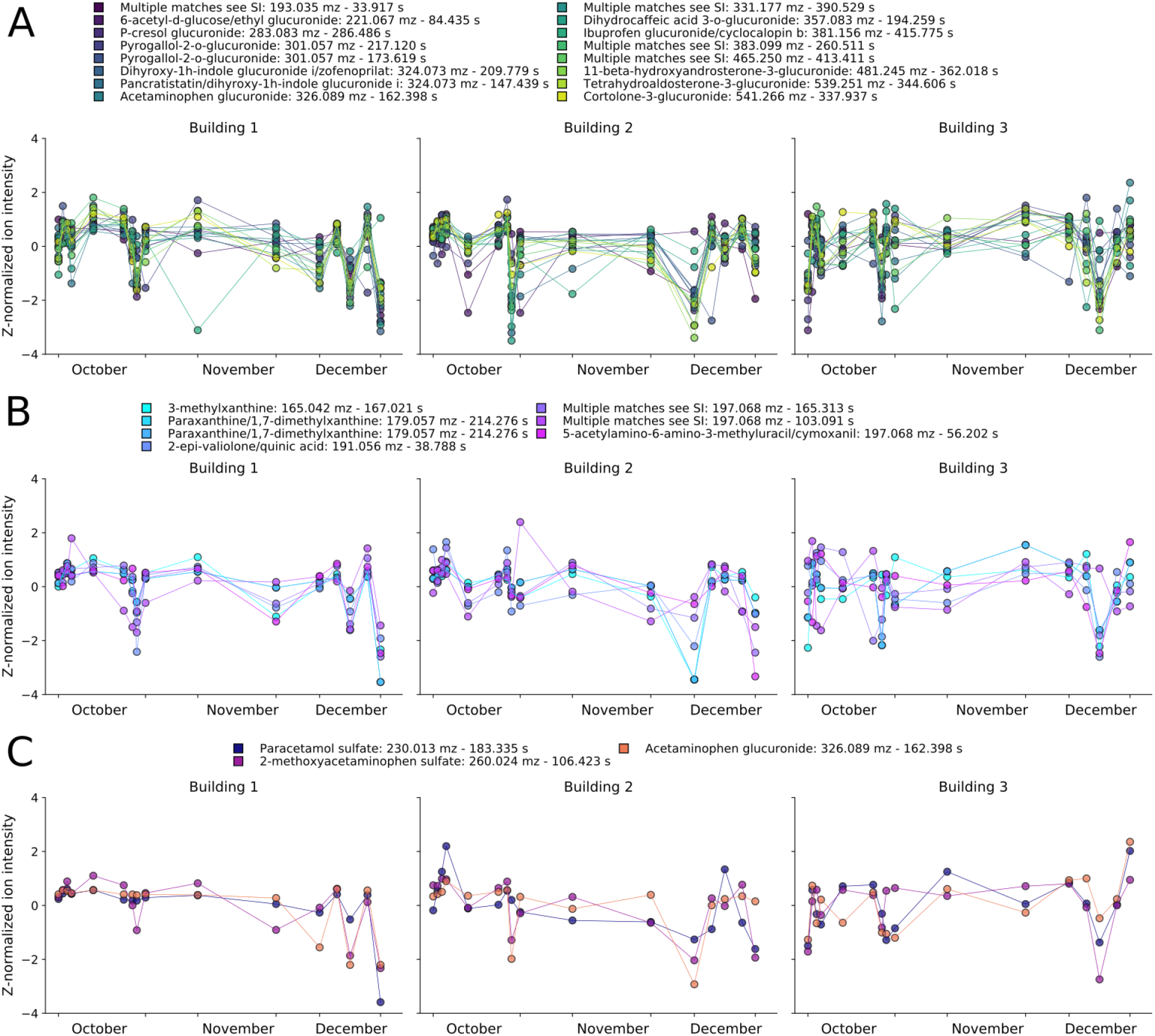
High temporal sampling allows for monitoring putative metabolite levels and dynamics in a building-specific manner. Z-normalized time courses of (A) glucuronide-related, (B) caffeine-related, and (C) acetaminophen-related putative metabolites across the three buildings. See Table S4 for all putative identifications.

### 3. Dense temporal data draws new feature correlations

Analyzing individual fTDs within and between buildings allows comparison of building-to-building similarity or lack thereof. To do so requires calculating feature similarities using through-time distance measurements. These measurements reveal feature correlations, something not necessarily possible with single time point measurements.

#### 3.1. Different buildings show few similar fTDs while many are correlated within a building

A large number of highly similar fTDs were observed internal to buildings but almost no similarities were observed between buildings. To study the similarity between z-normalized time series, we analysed a set of time series at different Euclidean distances (Figure 6A). This led us to set two similarity cutoff thresholds; 1.5 for high stringency and 2.82 for low stringency. A histogram of distances between all time series (an all-to-all comparison), displayed an increased number of similar time series within each building, while between buildings there were few similar time series up to a Euclidean distance of ~2.5 (Figure 6B). The high intrabuilding zero-distance bin primarily corresponded to distances calculated between a feature and itself, plus a small number of very similar features. For the different comparisons, the majority of similar time series belonged to pairs in which both features had high average intensities; few belonged to pairs in which one feature had low average intensity, possibly due to instrumental noise (Figure S3 and S4). This observation suggested that high similarity between fTDs did not arise from comparisons to normalized background or noise features.

**Figure 6.**
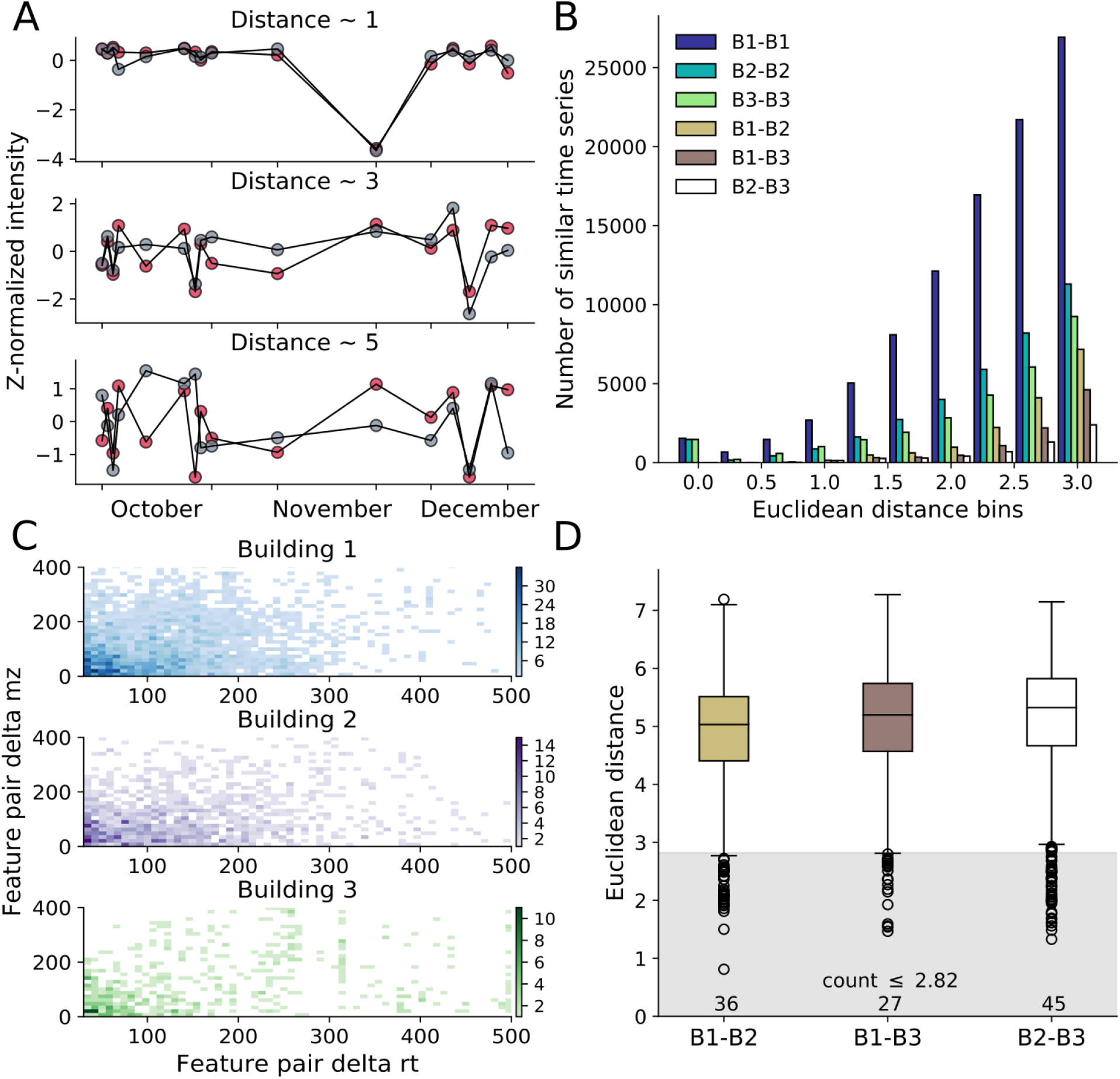
Many features display highly similar dynamics within, but not between buildings, with a large number being unrelated in mass and retention time. (A) Example distance plots for z-normalized data for distances of ~1, 3 and 5. (B) Histogram of all-to-all feature Euclidean distance calculations for different building pairs. (C) Within-building two dimensional histograms of all-to-all feature comparisons for which the Euclidean distance was < 1.5 and the difference in retention time was > 30 s, plotted versus the feature pair’s delta mz (y-axis). (D) Box plot of Euclidean distances between identical features across different buildings. The gray region holds all features below a distance of 2.82.

Further analysis on the large number of similar fTDs within a building, revealed that many of the feature pairs (at the high stringency cutoff) possessed large differences in mz and rt (Figure 6C, Figure S5A for complete range). A 2-D histogram of only those feature pairs that differed by at least 30 s in rt showed that many of the shared temporal dynamics differed by 30–100 s and 0–100 Da. A large number of feature pairs corresponded to mz and rt differences much greater than 100 s and 100 Da; overall, feature similarities were observed across the full mz and rt domains (Figure S5A). A comparable, but more populated, 2-D histogram was obtained with the low stringency similarity cutoff (Figure S5B and C).

The distance between a feature and its corresponding feature across buildings (a one-to-one comparison) demonstrated that only a few features (36, 27 or 45) possessed similar temporal dynamics between buildings, even at the more inclusive low stringency cutoff (Figure 6D). This analysis revealed that the majority of features displayed markedly different temporal dynamics, as the median distance was larger than five for each comparison.

### 4. Machine learning classifies buildings and finds building-specific features dynamics

Orthogonal to the approach of section 3, the complete feature profile of a single day provides a means for isolating community-specific feature information. Using an alternative objective—in this case, robust, across-day building classification—with simple machine learning models, it is possible to extract features with unique building-to-building temporal patterns.

#### 4.1. Single day feature profiles allow for building classification and find building-specific fTDs

L1-regularized logistic regression (L1-LR) and random forest (RF) models provided high classification performance and revealed building-differentiating features. We used a one-versus-the-rest approach for model training, for which each input corresponded to all feature values from a single day. Importantly, we used standardized log-transformed features intensities, not temporal z-normalized values. Receiver operating characteristic (ROC) area under the curve (AUC) analysis demonstrated high building classification AUC for all three comparisons with L1-LR models (0.934 ± 0.068 mean and standard deviation of 50-fold repeated model training, Figure 7A). The RF model performed slightly worse (AUC=0.893 ± 0.075, Figure S6), but provided additional insight from the set of features used.

**Figure 7.**
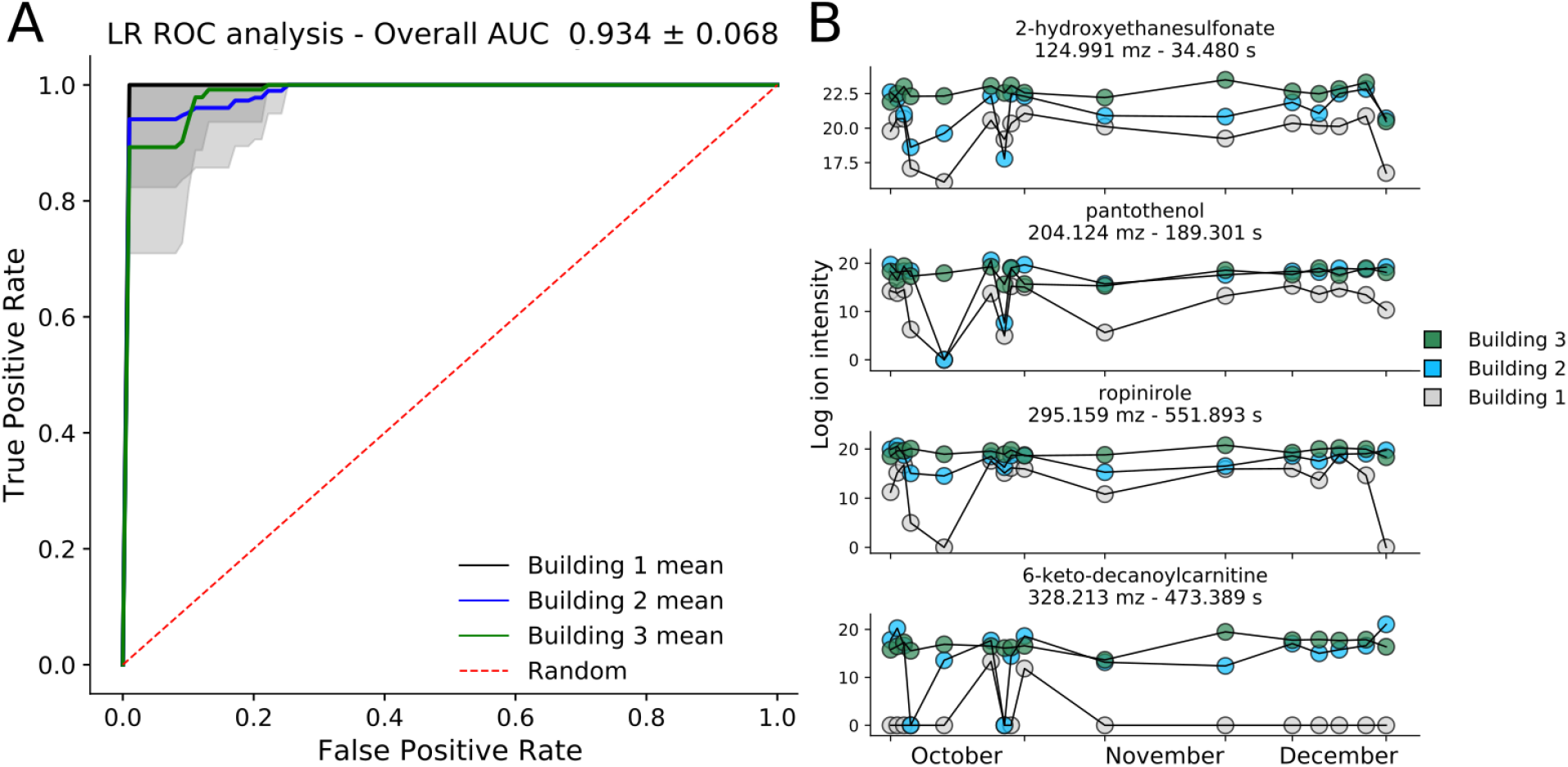
Single-day feature profiles allow accurate building classification and uncover features with building-specific fTDs. (A) ROC-AUC plot (true positive versus false positive rate) for the three one-versus-the-rest building comparisons, using individual days as data points with associated ion intensities as features. AUC plots and scores come from 50x repeated model training on randomized train-test data splits. (B) Putative identification (level 2) and log ion intensity time courses of four select, high-importance metabolites (see *Methods*) that contribute to model performance.

Features that were used in at least 40 of the independent L1-LR models and that possessed an average importance value greater than 0.005 across all 50 RF models demonstrated unique building dynamics. Four such feature time series are depicted in Figure 7B, of which three were highlighted by alternative methods (2-hydroxyethanesulfonate, pantothenol and ropinirole). The urine-related metabolite 2-hydroxyethanesulfonate and pantothenol both displayed lower ion intensity levels in the multipurpose-use Building 1 than in the residential buildings; similarly 6-keto-decanoylcarnitine, a urine metabolite used in non-muscle invasive bladder cancer diagnostic models^32^, was mostly absent in Building 1 but appeared at high levels in the other two (Figure 7B). Beyond these four, many of the important metabolites were either stable and statistically significant between buildings, or possessed alternative unstable metabolite class labels (Tables S5 and S6). This minimally biased data-driven modeling, largely recapitulated the findings of traditional statistical and temporal analyses, and discovered additional building-unique metabolite dynamics.

### 5. Grouping temporally similar features suggests targets for follow-up studies

We extracted additional features that were temporally related to the set of metabolites identified by our analyses. We grouped features temporally similar to each of the select metabolites for all three buildings, and analysed between-building, feature-pair cluster co-occurrences. This directly suggested features, and possibly hypotheses, for specific follow-up experiments, and may comment on shared controlling processes (chemical, biological, etc.) that govern feature dynamics.

#### 5.1 Intrabuilding temporal similarity and interbuilding co-clustering link possibly unrelated features

Analyzing the ‘important features’ (IFs, Tables S2-5) highlighted by our methods, we found that there were numerous, likely unrelated features within buildings that were highly correlated to the IFs. The IFs included the ML model features, putatively named metabolite groups, select unstable features, statistically significant stable features, and the urine, drug, sweetener, and cosmetics-related features. Metric multidimensional scaling (MDS) of the groups of features sharing fTDs with the IFs showed varying levels of clustering and co-clustering (Figure 8). Because many features were shared among multiple clusters, but only assigned to the largest, many groups displayed overlap in this two-dimensional space. Similar to other methods, MDS revealed that many features and putative metabolites with significantly different mz and rt values grouped with some of the most prominent IFs, many of which are believed to be human-related (Figure 8D). The clusters suggested unknown features that may originate from the same source, for which additional analysis is needed for identification.

**Figure 8.**
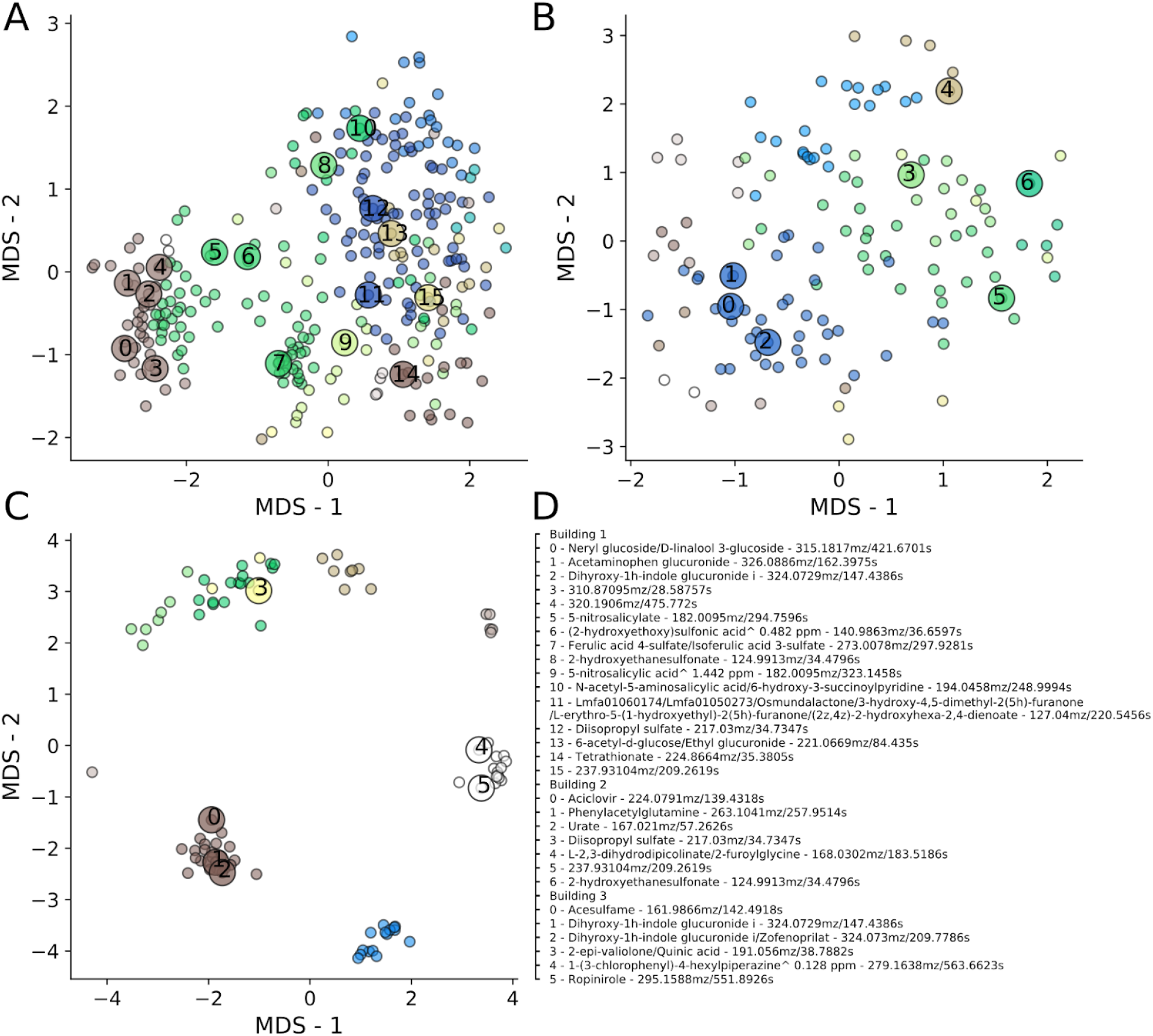
MDS analysis finds many putative metabolites or features that are temporally similar to select important features. MDS analysis of (A) Building 1, (B) Building 2, and (C) Building 3 feature groups with > 20 (A and B) or > 5 (C) similar features (as defined by intrabuilding feature-to-feature distance calculations). The ‘important features’ (IFs), to which distances from all other features were calculated, originated from the machine learning models, stability types, and other putatively named metabolites. Colors correspond to features that possess a Euclidean distance < 1.5 and group together. IFs are shown with numbers and large circles, and are labeled in (D). All names are level 2 identifications, unless ending with an ^ corresponding to level 3 identification with ppm error.

Between buildings we observed many co-clustered sets of features, despite most co-clustered features possessing different dynamics in each building. Using the intrabuilding K-means clusters, we found intersecting sets of features between pairs of buildings as well as between all three buildings (Figure S7). Again, this revealed groups of features with significantly different mz and rt values and that co-clustered in either two or all three buildings. The co-clustering of specific feature groups across buildings, despite building-specific temporal dynamics (Figure 6D), lends additional credence to the observed associations. Co-clustering across buildings, along with MDS analysis, highlighted numerous correlations between seemingly unrelated features, suggesting specific compounds or unidentified mz and rt pairs for follow-up analysis. Thus fTD analysis may further our understanding of specific chemicals (e.g. pesticides or drugs) by suggesting other small molecules that would otherwise appear unrelated, but that may have been introduced into the waste stream by similar controlling processes.

## Discussion

Previous temporal studies of wastewater metabolomics have examined seasonal variation33 or larger, aggregated populations at multiple time scales (often 24-hour aggregates), using downstream wastewater treatment plant (WWTP) sampling^16,34–36^. Here, we demonstrated the utility of short duration, high frequency (daily) direct building sampling to understand through-time statistical properties, individual feature temporal dynamics (fTDs), and clusters of temporally related features of single-building wastewater. Adding this temporal component, without significant wastewater aggregation, may benefit future wastewater metabolomics studies by revealing community-specific metabolite dynamics and the daily burden of environmental pollutants, drugs, or lifestyle-related small molecules. Longitudinal sampling uncovered highly dynamic and unique building profiles that would be lost with WWTP-based sampling^23^ or by sampling over longer time periods. Along with building-specific information, we presented a series of methodological techniques that provide orthogonal and overlapping information for the analysis of longitudinal untargeted wastewater metabolomics. We highlight several main findings: the importance of temporal data collection, the utility of untargeted metabolomics for community monitoring, data-driven methods for information extraction, and the importance of direct, building sampling.

Temporal data acquisition, in combination with clustered feature modeling and time series comparisons, provides information not available from stationary statistical properties alone. fTDs show that features may possess similar through-time mean intensities in different buildings, but with different temporal dynamics. These building-specific fluctuations may provide information on health-related events, given additional chemical identification. In light of the high level of feature spiking and subsequent rapid mean reversion, it is likely that this study was not sampled at a high enough frequency for much of its duration. As expected due to different individuals generating waste within and across days, autocorrelation drops rapidly (even with a time lag of one) for the five largest clusters in each building, indicating minimal information transfer between time points and in line with the results of the Gaussian process modeling (Figure S8). Frequent sampling may help explain select feature dynamics. For example, recreational drugs may be used at higher frequency on weekends rather than weekdays; thus, one might expect higher intensities on weekends. This suggests that future studies should sample daily and—in certain circumstances—multiple times per day, requiring additional device engineering for fully automated sampling, unlike the current manual process. Finally, the importance of short, high-frequency sampling can be seen in our theoretical waste stream mixing analysis, which suggested that much of the dynamics information is lost with the addition of only a few additional waste streams.

An unanticipated finding from this temporal analysis was that numerous features display highly similar temporal dynamics within buildings. While many of these feature pairs correspond to different adducts or isotopes, a large number appear to be attributable to different metabolites, as their masses and retention times can differ by hundreds of Daltons and seconds (Figures 6C, 8 and Figure S7). Specifically, focusing on highly correlated fTDs offers a means to discover new molecules (or features) linked to other molecules, events, or processes with known sets of features. Whether or not all of these features correspond to real metabolites, this information is readily supplied by fTDs and suggests perhaps non-obvious connections. In short, this approach may act as a hypothesis-generating method while also providing information about daily metabolite usage or exposure.

Untargeted metabolomics represents an information-rich method to monitor a community’s health and behavior. We putatively identified a host of human-related metabolites, most notably related to drugs, cosmetics, and food. For these, we observed that many features displayed similar intensities through time, but that their stability was frequently different across buildings. This unexpected observation of correlation between features’ mean intensities between buildings resulted in only a small handful being statistically significant. Yet this small number appeared to provide important, building-specific information (Figure 2C). Further, the buildings displayed a general trend in terms of overall intensity levels, with most intensities being higher in Building 2 than Building 1, and most in Building 1 being higher than those in Building 3 (Figure 2A)—likely reflecting the number of individuals using the toilet during the sampling period. While providing the potential for significant public health information, this method is limited by the ability to chemically identify each of the features observed. In addition, the use of only a single MS ionization mode and LC column type prevented a more complete report on the small molecule output of the buildings. These limitations warrant additional studies to expand feature-to-metabolite naming along with the use of select standards to validate putative metabolites.

A data-driven approach, based on classifying buildings using the features of a single day, recovers many of the features identified as important in other types of analysis, but also provides additional metabolites and features for follow up. Importantly, it extracts information relevant to each of the buildings in a minimally biased manner. The machine learning (ML) models we used found features in a manner complementary to the other presented methods, and demonstrate that it is possible to classify which building generated a specific, single-day waste profile. In addition to finding most of the 16 statistically significant features, the models also found features that belong to multiple classes of metabolite dynamics. For instance, 6-keto-decanoylcarnitine was important for building classification and was found to be unstable in Buildings 1 (class 1) and 2 (class 3), but stable in Building 3 (Tables S5 and S6). Our methods suggest a workflow for future temporal studies with the specific aim of public health monitoring. For instance, given data from well characterized control buildings and a new building of interest, using ML models and fTD clustering may help identify and track the dynamics of compounds in target communities.

Although direct building sampling was not the specific focus of this study, it was critical for this work, and our findings support the applicability of this technique for community-specific wastewater epidemiology. Most small-population studies have focused on targeted methods for measuring various drugs, with minimal temporal information^10,37^. Our recent 24-hour study likewise demonstrated the utility of upstream sampling, but of larger populations^23^. Single-building sampling minimizes the amount of time—and thus sample degradation—between sample generation and collection. This may address a potential source of uncertainty and error in WWTP-based measurements of population size or monitoring of illicit drug consumption^38–41^. Likewise, direct building sampling bypasses the issue of wastewater mixing or of occasional septage pumping into WWTPs, which may obfuscate fTDs or bias monitoring^1^. Thus, applications of the presented sampling method may include estimating population sizes and per capita feature values, and monitoring sporadic features.

High frequency, close-to-source sampling may, however, pose an ethical quandary. As the size of the population decreases, so does the anonymity of the results. For this reason, community-specific research must be conducted in such a way that personal health information remains confidential, and minimal negative consequences are experienced by the community under study^42^.

## Conclusion

Understanding and monitoring community health is a challenging but important problem for which longitudinal, untargeted metabolomics of single-building wastewater may prove beneficial. We demonstrate several methods to analyze such temporally resolved measurements, including through-time statistical properties, comparing feature temporal dynamics (fTDs) via Euclidean distance-based similarity metrics, and machine-learning modeling for feature extraction. Temporal analysis commonly shows that features undergo unpredictable and dramatic dynamics. We find that proximal buildings do not share common fTDs, while in contrast, a large number of similar features are found within each building. We putatively identify and track the dynamics of small molecules, compare them between buildings, and propose numerous links between features and metabolites that differ significantly in retention time and mass-to-charge ratio. These observations suggest that longitudinal studies, using daily sampling, can provide insight beyond the statistical averaging inherent to bulk wastewater sampling. We suggest that even higher frequency sampling, using additional analytical techniques, will only further improve our ability to provide important and actionable community-specific public health information.

## Methods

### Sample collection

Samples were collected from street-level manholes located outside of three buildings: one multipurpose-use building (Building 1), and two residential buildings (Buildings 2 and 3). We used a commercial peristaltic pump (Boxer) to continuously collect wastewater samples for 3 hours starting from 9:00 AM for Building 1 and 8:00 AM for Buildings 2 and 3. The peristaltic pump was programmed to pump wastewater at a rate of 5.55 mL/min over a 3-hour period into a 1 L polycarbonate bottle (Thermo Scientific) stored on ice, for total volume of 1 L of wastewater. 100 mL of each sample were then filtered separately through a 0.2 μM PTFE membrane filter (Millipore) using a glass filtration apparatus (Glassco) to remove bacteria and debris. All filtration glassware and polycarbonate bottles were acid washed with hydrochloric acid and autoclaved prior to filtration. The filtrate was collected in amber glass vials, the pH was adjusted to between 2 and 3, and stored at −80 □C, all in less than 2 hours post sampling.

### Liquid chromatography-mass spectrometry

10 μl of sample was analyzed via LCMS using a Vanquish ultra-performance liquid chromatography system coupled to an Orbitrap Fusion Lumos (Thermo Scientific) via a heated electrospray ionization (ESI) source. Data was collected in negative ionization mode with data-dependent secondary mass spectra (MS/MS) obtained via high-energy collisional dissociation (HCD, mass resolution 15,000 and collision energy of 35 arbitrary units, automatic gain control, AGT, of 5.0e4 and max injection time, IT, of 22 ms). The full MS resolution was 120,000 at 200 mz with an AGT target of 4.0e5 and a maximum IT of 50 ms. The quadrupole isolation width was set at 1.0 m/z. ESI was carried out at a source voltage of 2600 kV for negative ion mode with a capillary temperature of 350 □C, vaporizer temperature of 400 □C, and sheath, auxiliary, and sweep gases at 55, 20, and 1 arbitrary units, respectively.

Chromatographic separation was performed on a Waters Acquity HSS T3 column (2.1 × 100 mm, 1.8 μm) equipped with a Vanguard pre-column and maintained at 40 □C. The column was eluted with (A) 0.1% formic acid in water and (B) 0.1% formic acid in acetonitrile at a flow rate of 0.5 mL min^-1^.The gradient started at 1% B for 1 min, ramped to 15% B from 1–3 min, ramped to 50% from 3–6 min, ramped to 95% B from 6–9 min, held until 10 min, ramped to 1% from 10–10.2 min, and finally held at 1% B (total gradient time 12 min). Run order was randomized over two batches of samples with pooled quality control samples run intermittently (every 6 or 7 samples) along with MilliQ water blanks to account for the general background of solvent system and mass spectrometer.

### Data processing

Python 3.6.5 with scikit-learn version 0.19.1 as well as R 3.5.1 were used for processing and analysis. Following data acquisition, all data files were converted to an open source file format (.mzML) via a custom wrapper (msconvert_ee.py) of the program MSConvert in the ProteoWizard suite^43^. All files were then processed as a single batch with a custom python wrapper script (full_ipo_xcms.py) of both IPO^44^ and then subsequent XCMS^45^ processing. The parameters for XCMS were: CentWave (ppm=10, peakwidth=(5,15), snthresh=(100), prefilter=(4,10,000), mzCenterFun=wMean, integrate=2, mzdiff=-0.005, noise=50,000), ObiwarpParam (binsize=0.1, response=1, distFun=cor_opt, gapInit=0.3, gapExtend=2.4, factorDiag=2, factorGap=1), PeakDensityParam (bw=10, minFraction=0.05, minSamples=1, binSize=0.002, maxFeatures=50), mode (negative). In addition to aligning and extracting peak information, this program automatically extracted all MS/MS spectra and saved as a separate.mgf file for use in the metabolite naming pipeline.

All features were binary log transformed following the filling of empty values with one. The data was corrected for run order using the local (two closest run order-flanking quality control, QC, samples) and global (all) QC feature values where the normalized feature intensity was calculated with the following formula:

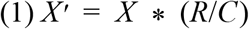

Here *X*’ corresponds to the corrected value, *X* the input feature value, *R* the global average of the feature over all QC sample and *C* the local feature QC average. All samples were corrected in this way.

Next, features with pooled sample coefficient of variance in excess of 0.3 were removed from further analysis. All samples were then blank subtracted (mean blank intensity for each feature) and resulting values less than 0 were set to 0. Finally, only features for which the cross building sum of log intensities was greater than 100 were kept as the remainder likely belonged to background signals.

### Putative metabolite identification

For ease of figure presentation when named features or lists thereof exceeded defined sizes, the name was replaced with ‘Name too long – See SI’ or ‘Many primary mz matches – See SI’ for which each mz-rt tuple can be mapped to names in the supplementary information and corresponding table. Minimum reporting standard level 2 corresponded to a feature’s secondary mass spectrum matched to an *in silico* fragmentation spectrum plus database mass match, level 3 was purely a database mass match and level 4 was unknown^24^. Given only putative identification throughout, all names should be interpreted with caution.

Identification was automated using custom python scripts, outlined in the supplementary information and associated Github pages. It performed a primary mass-to-charge look up of the exact mass accounting for multiple possible adducts and isotopes ([M]^-^, [M-H]^-^, [M+Cl^-^]^-^, [M-H-H_2_O]^-^, [2M-H]^-^, [M-2H+Na^+^]^-^, [M-2H+K^+^]^-^, [M+(1–3)^13^C-H]^-^) in four databases: MetaCyc^46^, the Human Metabolite DataBase (HMDB)^47^, the Chemical Entities of Biological Interest (ChEBI)^48^, and LIPID MAPS^49^. For this lookup, a parts per million (ppm) error tolerance of 10 was used, we did not consider names associated with ppm error values > 5 to be real and discounted them for all analysis, this represented an identification level of 3. Following this matching, the named features with associated ppm error and adducts or isotopes were ranked according to a heuristic order of likelihood of being observed (see SI for ordering). Given equally ranked adducts or isotopes, priority was given to the chemicals with the lowest ppm error. The second part of the naming was to perform *in silico* fragmentation matching using MetFrag^50,51^ on all ions with secondary mass spectra (MS2) that were extracted during the initial XCMS feature extraction. For this, individual MetFrag programs were run in parallel on MS2 scans (on up to three different MS2 for the precursor ion), for all of the following parent ion possibilities: [M]^-^, [M-H]^-^, [M+Cl^-^]^-^, [M-H-H_2_O]^-^, [2M-H]^-^, [M-2H+Na^+^]^-^, and [M-2H+K^+^]^-^. The MetFrag outputs for each parent ion were combined and ranked in order of MetFrag probability, these names were then matched to the primary mass-named metabolites with matches being given the highest ranking in terms of metabolite identification (level 2). Following naming, some putative labels were removed, specifically those with select elements, polymers or ‘R-groups’ (SI section 1 and Table S1).

### Statistical analysis

Multiple comparison tests of feature intensity differences between more than two buildings were done with the non-parametric Kruskal-Wallis test followed by Benjamini-Hochberg FDR correction (scipy.stats.kruskal, statsmodels.stats.multitest.multipletests). For stationary statistical significance an FDR corrected P-value of less than 0.00001 was used. Linear regression on mean feature values and the associated correlation constants were extracted using scipy.stats.linregress.

### Feature time series summary statistics

Through-time mean log intensity and standard deviation values were calculated for all features in each building individually. Features with a standard deviation ≥ 3 were labeled unstable, and stable otherwise. Given these labels, building overlap analysis was performed using venn3 and venn3_circles (matplotlib_venn) for which all building overlaps were retained. To determine feature color in Figure 1D, a feature was considered to be stable between the compared buildings if it was stable in either (colored blue).

To classify the unstable features, the following tests were performed. First, if any log ion intensity value exceeded the highest value observed in the stable metabolites the feature was grouped into class 2. Next, the median intensity for the time course was calculated, those < 5 were placed into class 1. The remaining features were labeled class 3 which corresponded to generally stable and medium to high intensities with the occasional, but significant downward intensity drops.

### Temporal feature profile analysis—building clustering

All features were z-normalized through time (temporal mean subtracted and divided by the standard deviation) and each building was clustered by K-means clustering (sklearn.cluster.KMeans) using 100 clusters with all other parameters kept to their default values. Each feature was mapped to its corresponding mz and rt value for plotting as well as being grouped by cluster size. Mean cluster values were extracted for each of the clusters and used to train a GP regression model (sklearn.gaussian_process.GaussianProcessRegressor, alpha=0.0001) using a radial basis function kernel with a 0.125 length_scale parameter mixed linearly with a constant kernel with noise corresponding to the mean standard deviation of elements of the cluster (sklearn.gaussian_process.kernels.RBF and .ConstantKernel respectively).

### Waste stream simulations and mixing analysis

To match experimental data as accurately as possible, for each waste stream simulation we sampled 100 cluster means for the 16 days using a Cauchy distribution (scipy.stats.cauchy, loc=0, scale=0.3, size=1 and resampled if a value greater than 3.5 was obtained). We then sampled the number of intensity spikes (positive and negative) for the new wastestream from a Gaussian distribution (random.gauss, mean=0, standard deviation=1) and took the absolute value of the integer representation. We then randomly determined which days would have the spikes and the number of cluster across which this spike would occur (they normally occured over many clusters in one day) using 20% of the number of clusters (20) times the absolute, integer value of a Gaussian of 0 mean and unit standard deviation. The exact clusters with the spiking dynamics were probabilistically determined for each day independently using a decreasing exponential probability (0.002+e^-i^, where i is the cluster rank), the clusters were looped over (from largest to smallest) with a value in [0.0, 1) randomly chosen, if the value was less cluster’s probability threshold it was included in the spiking cluster. This was repeated until the total number of spiking clusters was satisfied. For these linked clusters, the mean values were resampled from the original cauchy distribution but forced to be between 1.75 and 3.

Many of the non-spiking feature intensities were similarly conserved across clusters. For this we sampled the number of days with ‘correlated’ features using the absolute integer value of a Gaussian with 0 mean and 3 for its standard deviation. The number of correlated clusters was performed as before but with a multiplicative factor of 70%; days of correlation were then probabilistically chosen according with 0.002+0.8e^-0.8i^ representing a cluster’s probability being included (this spreads the probability out over more clusters, especially to the smaller ones). For these clusters, new means were chosen from the original cauchy distribution but forced into the range of being less than 1.75.

Finally, once all cluster means were sampled, 1440 features were drawn (with cluster sizes proportional to the average cluster size in the three real buildings) using a Gaussian distribution with the cluster mean for each cluster and a scale of 0.3. All values were forced to be less than 3.2 as intensities of this magnitude were rarely if ever observed in the three real data sets.

For mixing, this waste stream profile generation process was repeated a select number of times. For all numbers of mixed waste streams, the three real buildings were mixed at random percentages with the simulated data to create a single mixed data set which was then clustered with K-means (100 clusters), and each cluster fit with a GP. The calculation of the all cluster mean to all cluster mean distance was performed using the distance_matrix function and then summed. To get statistical parameters for this process, the analysis was repeated 10 times to calculate a mean and standard deviation for each number of waste streams mixed.

### Temporal feature profile analysis—inter and intra building feature analysis

Time series distances were calculated via a Euclidean distance metric (scipy.spatial.distance.euclidean). All-to-all feature distance matrix calculation within and between buildings was performed using scipy.spatial.distance_matrix while the one-to-one distance of each feature to the corresponding feature in the other building using the euclidean function. For the all-to-all, feature pairs that met the similarity cutoff of either 1.5 or 2.82 were kept and further analyzed for both rt and mz difference between features.

### Machine learning model training and analysis

L1-LR and RF models were built to predict which building a single day-feature profile belonged to (sklearn.ensemble.RandomForestClassifier, and sklearn.linear_model.LogisticRegressionCV with the ‘saga’ solver). To gain statistics on model performance, each was trained 50 times on random, full data shuffles (sklearn.utils.shuffle). For each data shuffle, the days were randomly split 75:25 into train and test splits respectively. The log ion intensity data training split was standardized (sklearn.preprocessing.StandardScaler), and the test data transformed. Internal cross validation (3-fold) on the training split was performed internal to the LogisticRegressionCV class while the number of trees for the RF was set to 1000 requiring no cross validation. Following training, all models were evaluated on the fully held out test set. ROC-AUC analysis was performed for each separate model (using sklearn.metrics.roc_curve and .auc) during the testing phase. For the 50 models built, the feature coefficients of the L1-LR models or the feature importances from the RF were averaged. To isolate unique building features, averaged features that had non-zero feature coefficients in at least 40 of the L1-LR models and an averaged feature importance in the RF models of > 0.005 were extracted.

### ‘Important’ feature and cross building analysis

Temporal similarity values were calculated between all features to the IFs (all features from Tables S2-5) in each building separately using a Euclidean distance metric. Many features possessed a distance < 1.5 to multiple IFs and were counted for the size of each IF’s group. After calculating the size of each group, all features were then only assigned to the largest group they belonged to and those with sizes greater than 20 for Building 1 and 2 or 5 for Building 3 were input to metric MDS (sklearn.manifold.MDS).

To find features that co-clustered between buildings, the cluster memberships for all buildings were fully compared, including between all three buildings. If the set of features in the intersection of either 2 or 3 clusters (each from different buildings) was > 5, the cluster pair overlap of features was kept. A cluster pairing was only further analyzed if the difference in minimum and maximum rt was greater than 30 s.

## Data Availability

Packaged data set files along with XCMS processed data tables (.csv files) will be deposited in Zenodo upon publication.

## Code Availability

All code (.sh, .py, .R and .ipynb) to replicate the results of this paper will be on the following Github repositories upon publication: https://github.com/ethanev/Metabolite_lookup and https://github.com/ethanev/temporal_wastewater

## Acknowledgements

We acknowledge Dr. Melissa Kido Soule and Dr. Elizabeth Kujawinski and the funding sources of the WHOI FT-MS Users’ Facility (National Science Foundation MRI program OCE-0619608 and the Gordon and Betty Moore Foundation GMBF1214) for assistance with MS data acquisition. Funding was provided by the MIT Center for Microbiome Informatics & Therapeutics (CMIT) as well as the Underworlds project that is funded by the Kuwait-MIT Center for Natural Resources and the Environment along with the MIT Senseable City Lab.

## Author contributions

EDE, and EJA analyzed, interpreted data, and wrote the manuscript. CD, SI and SP performed sampling. All authors edited and provided feedback for the manuscript.

