## Supplementary Information for "Longitudinal wastewater sampling in buildings reveals temporal dynamics of metabolites"

**This file includes:**

1. Additional data processing methods
2. Additional data
3. References

### 1. Additional data processing methods

The following is an outline of how to process the data (similar information is on the associated Github pages), extract the primary feature table and associated secondary mass spectra (MS2) and then name the features from this XCMS output, for use in downstream analysis.

*File conversion*

To convert the file to .mzML format the following command was used on a window operating system, this was run in the directory that contained the mscovert.exe program, otherwise the script can be modified to accept an input msconvert.exe path:

```
.\path\to\msconvert_ee.py .\path\to\data
```

*XCMS processing*

To process the .mzML data, the following command line script is a template to run the full_ipo_xcms.py script that wraps both IPO and XCMS:

```
python full_ipo_xcms.py --data_type 'mzML' \
--in_path='/path/to/data/' \
# if IPO out files exist, otherwise remove the following line (ie for the first time running):
--ipo_files='IPO_1.out IPO_2.out IPO_3.out' \
--data_file='name_of_file_with_path_and_names_to_all_mzMLs.txt' \
--acq_mode='negative' \
--csv_out='out_file_name.csv' \
--out_path='/path/where/you/want/the/data/' \
--log_file='out_file_log_name.log'
```

This generated both the .csv file with features and their intensities as well as the .mgf file or metabolite naming.

*Feature naming*

For naming we note that we attempted to include possible isotopes, primarily multiple C13 peaks; however, we find these labels to not be realistic as most of the time only the C13 version of a compound would be found and not the C12, this makes little sense chemically and thus we urge the user to not pay attention to these labels and remove them. Additionally, we have found 'hydrate' compounds as well as 'sodium' or 'potassium' adducts in some names, these too should be ignored.

1) Use parsing_metabolite_dbs.py

This will read in and process the various databases (HMDB[2], MetaCyc[3], ChEBI[4] and LIPID MAPS[5]: hmdb_metabolites.xml, compounds.dat, ChEBI_complete.sdf and structures.sdf) but you must run each individually with the appropriate command line flag after downloading the appropriate databases, these are not included in our data or github page.

2) Use mapping_mz_to_metabolites.py
This will perform the mapping from mz features to small molecule names in the databases. It makes a csv file for each dataset and outputs them to a user specified directory. Example command line (all options are detailed in the script itself and besides path, this command was used for this manuscript):

Python ./mapping_mz_to_metabolites.py -f
'./path/in_file_after_ipython_feat_processing.csv' -p 10 -c 1 -m 'negative' -o
'./path/out_file.csv'

3) Use metabolite_calling.py
This will go through the csv file from the previous step, cluster by defined retention time (rt) for peak grouping, find similar chemicals and call the best (if possible) metabolite. The metabolite calling or 'ranking' had the following priorities: $[M\text{-}H]^-$ > $[M\text{+}Cl]^-$ = $[M\text{-}H\text{-}H_2O]^-$ = $[M]^-$ > $[2M\text{-}H]^-$ = $[M\text{-}2H\text{+}Na^+]^-$ = $[M\text{-}2H\text{+}K^+]^-$ = $[M\text{+}(1\text{-}3)^{13}C\text{-}H]^-$. This creates a file with the following general name: all_voted_in_file_from_step_2.csv An example command line is:
python ./metabolite_calling.py -t 5 -r 10 -f "./path/in_file_from_step_2.csv" -s ',' -o './path/'

To get MS-MS verification:
1) run parse_mgf_comb_feat_prep_metfrag.py on the all_voted_* file from the voting above. This must be run in the same folder as the Metfrag program[6]. It runs in parallel all the metfrag programs generated for 3 (or fewer if that's all there are) MS2 spectra for a single mz/rt pair for each possible adduct. This requires you to have the extra database files: hmdb_2017-07-23.csv, kegg_2017-07-23.csv, lipidmaps.csv (not provided since we do not own this data). Also, **it requires a subfolder named msms_out**, this is where all the output files will be placed. Example command line:
python parse_mgf_comb_feat_prep_metfrag.py -f './all_voted_in_file_from_step_2.csv' -c 2 -r 3 -m './mgf_file_from_xcms.mgf' -v 'true'

2) run combine_metfrag_w_votes.py
This will combine the output of the metfrag program (lots of individual files with the voted results) to find the best metabolite names. Examples command line:
python combine_metfrag_w_votes.py -f './all_voted_in_file_from_step_2.csv' -c 2 -r 3 -m './path/to/msms_out/' -v 'true' -n 'True' -o './final_named_metabolites_ordered.csv'

3) To match the named metabolites to the features and their intensities use the data_processing.ipynb (along with its primary function of finding the best features to work with) and follow all the way to the end and save the final output file 'partly_cleaned_metals_combined_mz_rt_metfrag_votes_isotopes_mzrt_named.csv'. This

remove most implausible compounds according to Table S1.

To perform data analysis and reproduce results of the papers follow the four following notebooks with save = True and processed = False for the first time through and then reversed so as not to retrain the various models. Note, results may differ due to different random seed for algorithms. Use the following workflow:
1) data_processing.ipynb
2) splitting_stable_unstable_metabs_pub.ipynb
3) Building_and_day_classification.ipynb
4) temporal_dynamics_analysis.ipynb

**Table S1. Name components that caused a label to be discarded.** If any of the following showed up in the name or chemical formula for a putative name for a feature, it was removed from consideration as a plausible name.

| | |
|---|---|
| **Elements** | 'Ac','Mt','Ag','Al','Na','Am','Nb','Ar','Nd','As','Ne','At','Au','Ba','Be','Og', 'Bi','Pa','Pb','Ca','Pd','Cd','Pm','Ce','Pr','Pt','Cm','Pu','Ra','Rb','Cr','Re','Rf', 'Cu','Rg','Rh','Db','Rn','Ds','Ru','Dy','Er','Sb','Es','Eu','Se','Sg','Fe','Si','Fl', 'Sm','Fm','Fr','Sr','Ga','T','Gd','Ta','Ge','Tb','Tc','He','Te','Th','Hg','Ti','Tl', 'Tm','Ts','U','Ir','V','K','W','Kr','Xe','La','Y','Li','Yb','Lr','Zn','Lu','Zr','Lv', 'Mc','Md',’Mg','Mn','Mo', 'R', 'X’ |
| **Group_names** | 'Group','hyltin','stann','nickel','medronic','copolymer','residue', 'Bioconjugate','conjugate','radical','borthiin','hydrochloride','vanadyl', 'intermediate', 'all-trans-retinol-R' |

### 2. Additional Data

**Table S2. mz, rt and the best name for each putative metabolite of special interest.** **denotes level 3 ID, otherwise, if named, it is a level 2 ID. For level 2 IDs only the chemical name and MetFrag probability are presented, for level 3 there is the name, ppm error, adduct and database matched and chemical formula.

| mz | rt | metfrag_matched_best_guess |
|---|---|---|
| 124.99134 | 34.47959 | 2-hydroxyethanesulfonate, 1.0 |
| 168.0302 | 183.51859 | l-2,3-dihydrodipicolinate, 1.0, **2-furoylglycine**, 1.0 |
| 181.99179 | 206.15887 | saccharin, 1.0 |
| 194.04581 | 248.9994 | n-acetyl-5-aminosalicylic acid, 1.0, 6-hydroxy-3-succinoylpyridine, 1.0 |
| 215.03296 | 34.81924 | 2-c-methyl-d-erythritol 4-phosphate, 1.0 |
| 254.91726 | 29.39754 | trichlorfon**, 6.002, M-H_chebi, C4H8Cl3O4P1 |
| 265.09139 | 285.8143 | alpha-bradyrhizose**, 5.013, M-H_chebi, C10H18O8 |
| 273.14599 | 239.91284 | 273.14599 mz /239.91284 s** - far too many compounds, not identifiable |
| 310.87095 | 28.58757 | 310.87095 mz /28.58757 s |
| 329.23383 | 396.91011 | 9,12,13-trihome(10), 1.0 |
| 331.22431 | 330.17615 | 331.22431 mz /330.17615 s |
| 362.94146 | 29.20236 | 362.94146 mz /29.20236 s |
| 387.28701 | 426.32416 | Methyl 2-(10-heptadecenyl)-6-hydroxybenzoate**, 8.908, M-H_hmdb, C25H40O3 |
| 405.14417 | 236.14125 | 2-[[2-[2-(2,3-dihydroindol-1-yl)-2-oxoethyl]-1-oxo-5-isoquinolinyl]oxy]acetic acid ethyl ester**, 3.826, M-H_chebi, C23H22N2O5 |
| 421.06048 | 384.55247 | N'-(3-chloro-2,4,6-trifluorophenyl)sulfonyl-1-adamantanecarbohydrazide**, 0.578, M-H_chebi, C17H18Cl1F3N2O3S1 |
| 446.90657 | 29.18975 | 446.90657 mz /29.18975 s |

**Table S3. mz, rt and the best name for each putative metabolite of mapping to temporal dynamics.** All IDs are level 2 with name and MetFrag probability.

| mz | rt | metfrag_matched_best_guess |
|---|---|---|
| 127.03997 | 220.54562 | lmfa01060174, 1.0, lmfa01050273, 1.0, osmundalactone, 1.0, 3-hydroxy-4,5-dimethyl-2(5h)-furanone, 1.0, l-erythro-5-(1-hydroxyethyl)-2(5h)-furanone, 1.0, (2z,4z)-2-hydroxyhexa-2,4-dienoate, 1.0 |
| 177.0403 | 38.5381 | l-gulonolactone, 1.0, 3,6-anhydro-l-galactonate, 1.0 |
| 157.03657 | 36.05959 | allantoin, 1.0 |
| 448.30749 | 445.62485 | glycodeoxycholic acid, 1.0, deoxycholic acid glycine conjugate, 1.0, chenodeoxyglycocholic acid, 1.0, glycodeoxycholate, 1.0, glycochenodeoxycholate, 1.0 |
| 212.00243 | 248.71144 | indoxyl sulfate, 1.0 |
| 230.14001 | 278.12369 | O-isobutyryl-l-carnitine, 1.0, o-butanoylcarnitine', 1.0 |

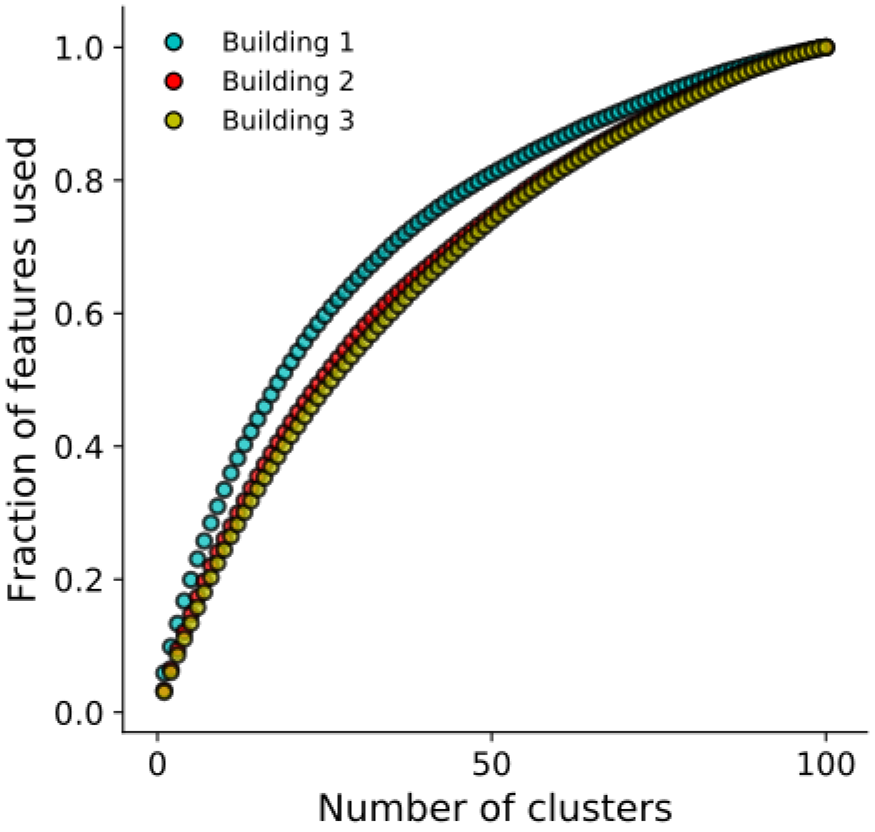


**Figure S1. Enrichment of features used relative to the number of clusters.** The clusters were first sorted by size and then for each number of clusters, the number of cluster members was summed for that value of combined clusters.

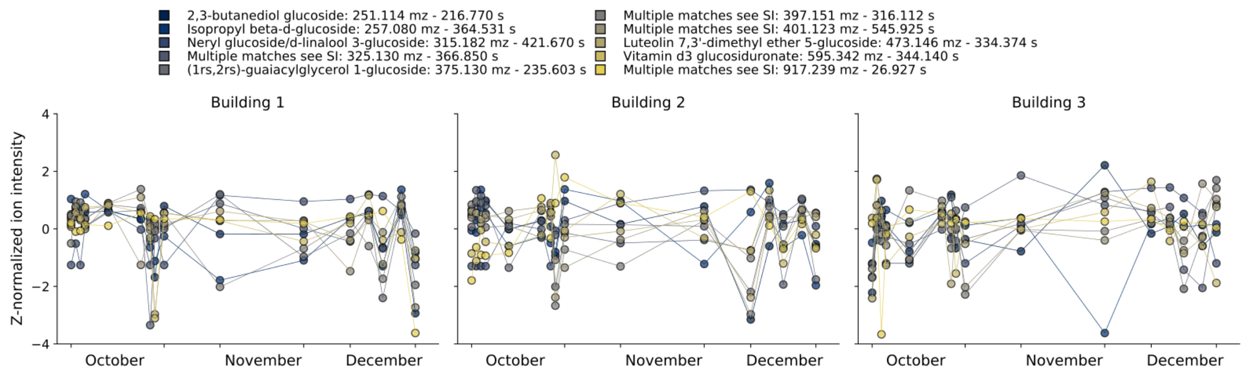


**Figure S2. Feature dynamics for possible glucoside-related metabolites.** Compounds with ‘glucosid’ in their name, whether matched at level 2 or 3 were included in the plot. See Table S4 for complete naming for those labeled as ‘Multiple matches see SI’.

**Table S4. mz, rt and putative names for select metabolite classes.** **denotes level 3 ID, otherwise if a name is supplied it is a level 2 ID. For level 2 IDs only the chemical name and MetFrag probability are presents, for level 3 there is the name, ppm error, adduct and database matched and chemical formula.

| mz | rt | metfrag_matched_best_guess |
|---|---|---|
| 193.03522 | 33.91739 | d-glucuronic acid', 1.0, 5-dehydro-l-gluconate, 1.0, duronic acid, 1.0, 5-keto-d-gluconate, 1.0 |
| 221.06687 | 84.43501 | 6-acetyl-d-glucose, 1.0, ethyl glucuronide, 1.0 |
| 283.08266 | 286.48602 | p-cresol glucuronide, 1.0 |
| 301.0569 | 217.11952 | pyrogallol-2-o-glucuronide, 1.0 |
| 301.05694 | 173.61856 | pyrogallol-2-o-glucuronide', 1.0 |
| 324.07296 | 209.77855 | dihyroxy-1h-indole glucuronide i, 1.0, zofenoprilat, 1.0 |
| 324.07294 | 147.43857 | pancratistatin, 1.0, dihyroxy-1h-indole glucuronide i, 1.0 |
| 326.08858 | 162.39754 | acetaminophen glucuronide, 1.0 |
| 331.17672 | 390.52876 | (2s,3s)-2-hydroxytridecane-1,2,3-tricarboxylic acid, 1.0, lmfa13010036, 1.0, (2s,3s)-2-hydroxytridecane-1,2,3-tricarboxylate, 1.0, neomenthol-glucuronide, 1.0 |
| 357.08321 | 194.25854 | dihydrocaffeic acid 3-o-glucuronide, 1.0 |
| 381.15606 | 415.77499 | ibuprofen glucuronide, 1.0, cyclocalopin b, 1.0 |
| 383.09898 | 260.51053 | 5-(3',5'-dihydroxyphenyl)-gamma-valerolactone 3-o-glucuronide, 1.0, 5-(3',4'-dihydroxyphenyl)-gamma-valerolactone-4'-o-glucuronide, 1.0 |
| 465.25007 | 413.41064 | etiocholan-3alpha-ol-17-one 3-glucuronide, 1.0, 3-alpha-hydroxy-5-alpha-androstane-17-one 3-d-glucuronide, 1.0 |
| 481.24499 | 362.0177 | 11-beta-hydroxyandrosterone-3-glucuronide, 1.0 |
| 539.25052 | 344.60582 | tetrahydroaldosterone-3-glucuronide, 1.0 |
| 541.26611 | 337.93658 | cortolone-3-glucuronide, 1.0 |
| 165.04177 | 167.02075 | 3-methylxanthine, 1.0 |
| 179.05748 | 214.27583 | paraxanthine, 1.0 |
| 191.05603 | 38.78821 | 2-epi-valiolone, 1.0, quinic acid, 1.0 |
| 197.068 | 165.31317 | 6-amino-5[n-methylformylamino]-1-methyluracil, 1.0, 5-acetylamino-6-amino-3-methyluracil, 1.0 |
| 197.06795 | 103.09128 | 5-acetylamino-6-amino-3-methyluracil, 1.0, 6-amino-5[n-methylformylamino]-1-methyluracil, 1.0 |
| 197.06798 | 56.20195 | 5-acetylamino-6-amino-3-methyluracil, 1.0, cymoxanil, 1.0 |
| 230.01308 | 183.33546 | paracetamol sulfate, 1.0 |
| 260.02376 | 106.42347 | 2-methoxyacetaminophen sulfate, 1.0 |
| 251.11394 | 216.76981 | 2,3-butanediol glucoside, 1.0 |
| 257.0798 | 364.5307 | isopropyl beta-d-glucoside, 1.0 |
| 315.18174 | 421.6701 | neryl glucoside, 1.0, d-linalool 3-glucoside, 1.0 |
| 325.12969 | 366.84988 | (3R,4R)-4,8-dihydroxy-3-((R)-2-hydroxypentyl)-6,7-dimethoxyisochroman-1-one**, 0.104, M-H_chebi, C16H22O7, Hinokitiol glucoside**, 0.104, M-H_chebi, C16H22O7 |
| 375.13023 | 235.60283 | (1rs,2rs)-guaiacylglycerol 1-glucoside, 1.0 |
| 397.15097 | 316.11212 | methyl (3x,10r)-dihydroxy-11-dodecene-6,8-diynoate 10-glucoside, 1.0, methyl 3,4-dihydroxy-5-prenylbenzoate 3-glucoside, 1.0 |
| 401.12314 | 545.92508 | 7-hydroxyflavanone 7-O-beta-D-glucoside**, 1.452, M-H_chebi, C21H22O8, (3'R,4'R)-3'-Epoxyangeloyloxy-4'-acetoxy-3',4'-dihydroseselin**, 1.452, M-H_chebi, C21H22O8, nobiletin**, 1.452, M-H_chebi, C21H22O8, graminone B**, 1.452, M-H_chebi, C21H22O8, 2-(2,5-dimethoxyphenyl)-5,6,7,8-tetramethoxy-4H-1-benzopyran-4-one**, 1.452, M-H_chebi, C21H22O8, 2-(3,5-dimethoxyphenyl)-5,6,7,8-tetramethoxy-4H-1-benzopyran-4-one**, 1.452, M-H_chebi, C21H22O8, 2-(3,4-dimethoxyphenyl)-3,5,6,7-tetramethoxy-1-benzopyran-4-one**, 1.452, M-H_chebi, C21H22O8, 5-[(5-methoxycarbonyl-2-methyl-3-furanyl)methoxy]-2-methyl-3-benzofurancarboxylic acid 2-methoxyethyl ester**, 1.452, M-H_chebi, C21H22O8, 5,6,7-trimethoxy-3-(3,4,5-trimethoxyphenyl)-1-benzopyran-4-one** , 1.452, M-H_chebi, C21H22O8 |
| 473.14596 | 334.37358 | luteolin 7,3'-dimethyl ether 5-glucoside, 1.0 |
| 595.34175 | 344.13965 | vitamin d3 glucosiduronate, 1.0 |
| 917.23945 | 26.92723 | Kaempferol 3-O-[6-(4-coumaroyl)-beta-D-glucosyl-(1->2)-beta-D-glucosyl-(1->2)-beta-D-glucoside]**, 4.058, M-H_chebi, C42H46O23, quercetin 3-O-alpha-L-[6'''-p-coumaroyl-beta-D-glucopyranosyl-(1->2)-rhamnopyranoside]-7-O-beta-D-glucopyranoside**, 4.058, M-H_chebi, C42H46O23 |

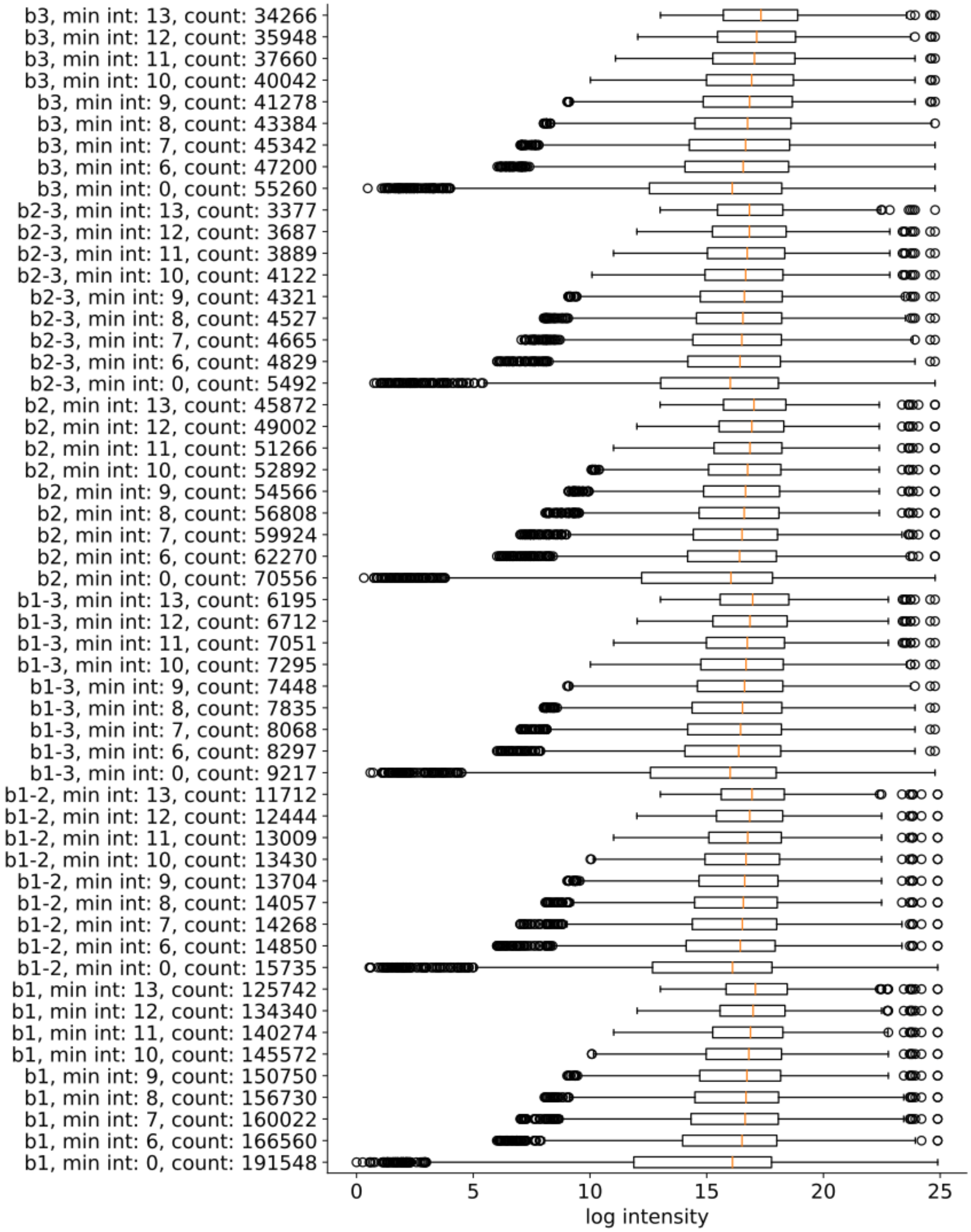


**Figure S3. Removing all temporally related feature pairs with at least one possessing a through-time mean intensity less than minimum intensity value (distance cutoff = 2.82 for similarity).**

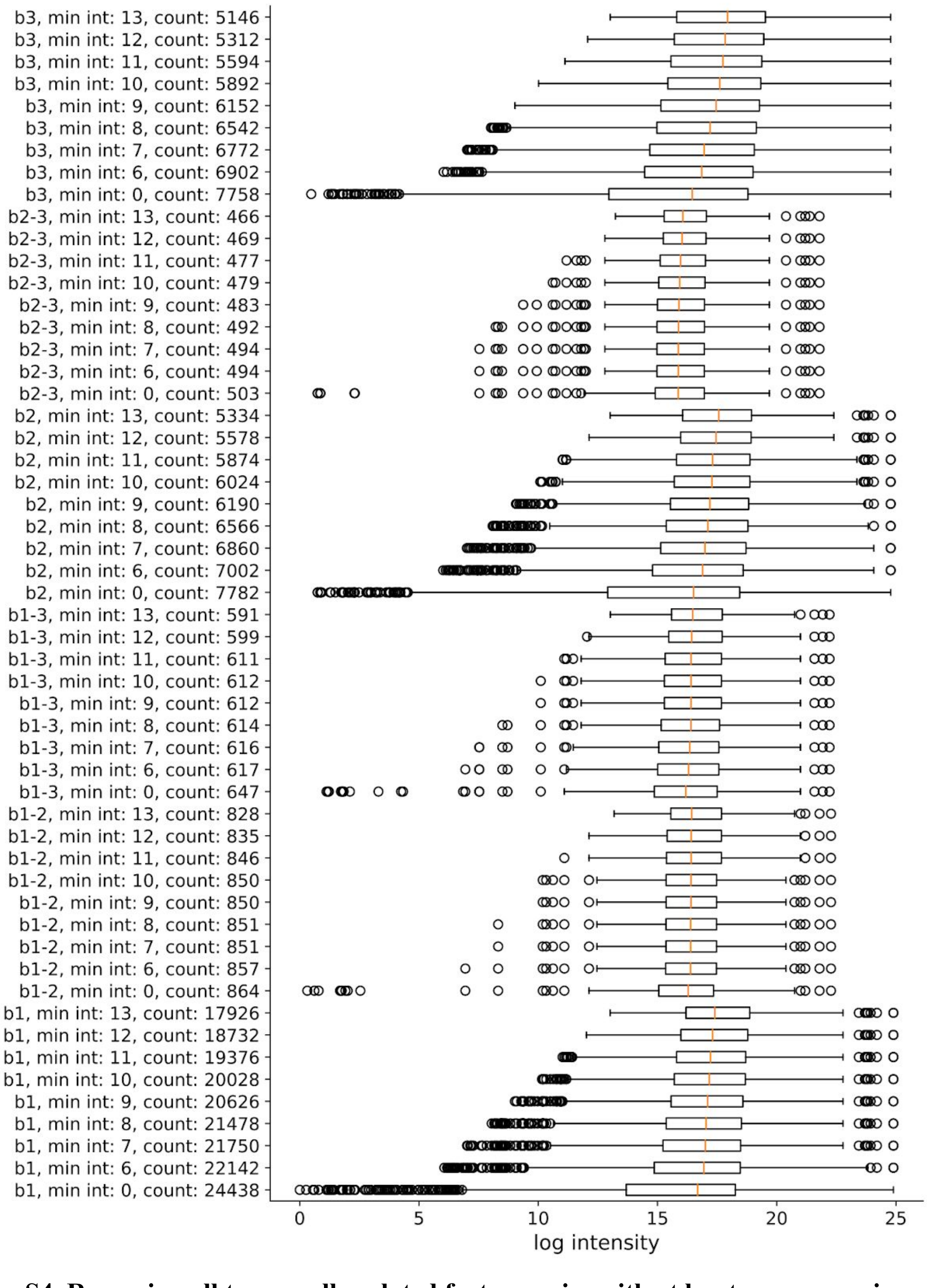


**Figure S4. Removing all temporally related feature pairs with at least one possessing a through-time mean intensity less than minimum intensity value (distance cutoff = 1.5 for similarity).**

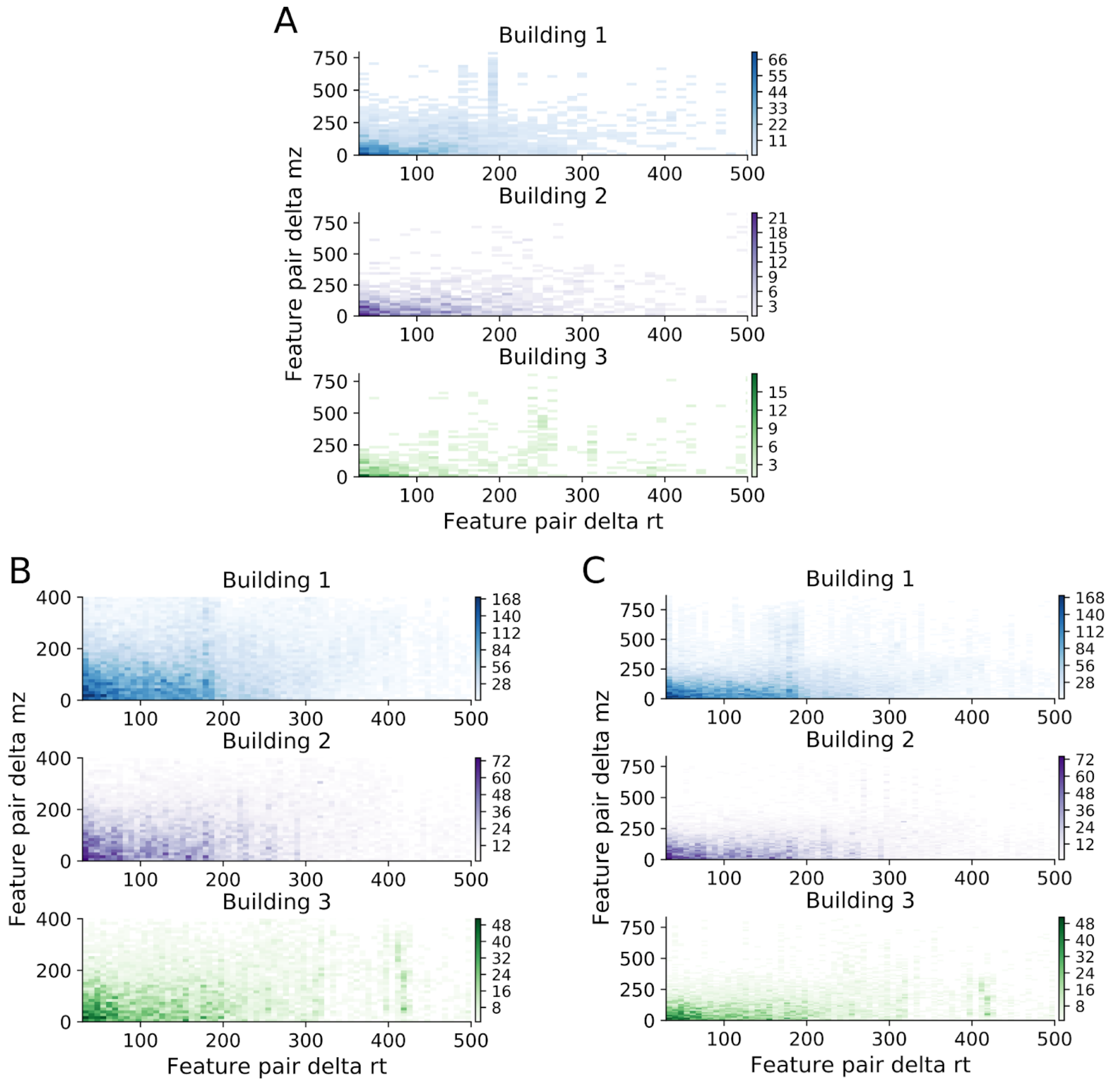


**Figure S5. Complete delta mz, rt 2-D histogram of temporally similar features for the three buildings at different distances.** (A) Full mz and rt domain for < 1.5 distance features. (B) Reduced and (C) full domain histograms for features with similarity < 2.82 in Euclidean distance.

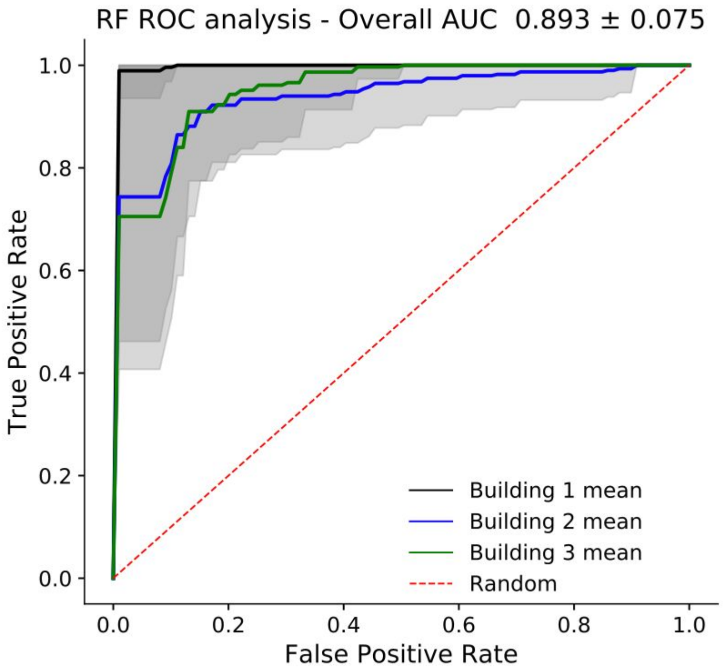


**Figure S6. Receiver operating characteristic (ROC) area under the curve (AUC) analysis for the random forest (RF) models.** Mean of the three AUCs presented as the overall AUC.

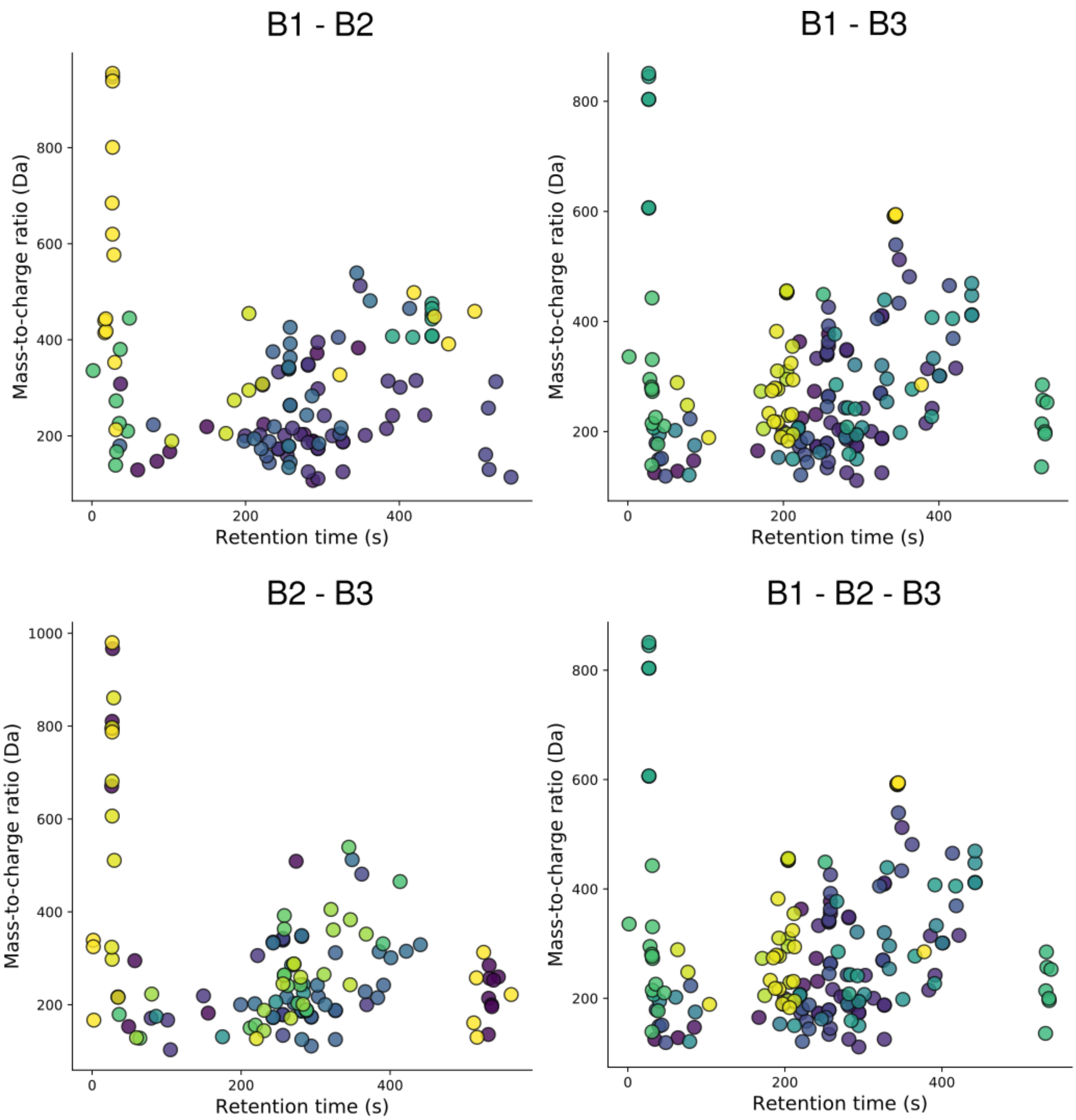


**Figure S7. Co-clustered features between buildings with at least 5 features shared in the clusters between the buildings and a minimum delta intracluster rt min and max of 30 s.** Each color corresponds to a commonly clustered features between the compared buildings.

**Table S5. Features important from both the RF and LR models.** Features with '**' denote that they are from solely primary mz database name matching (minimum reporting standards level 3) and where possible show a name, ppm error, which database and adduct was used and chemical formula. Features with just mz/rt with '**' indicate that there were far too many names for analysis. Features with solely a name and subsequent number are from mz-to-database match along with Metfrag identification.

| mz | rt | metfrag_matched_best_guess |
|---|---|---|
| 124.9913 | 34.4796 | 2-hydroxyethanesulfonate, 1.0 |
| 140.9863 | 36.6597 | (2-hydroxyethoxy)sulfonic acid**, 0.482, M-H_hmdb, C2H6O5S1 |
| 168.0302 | 183.5186 | l-2,3-dihydrodipicolinate, 1.0, 2-furoylglycine, 1.0 |
| 182.0095 | 323.1458 | 5-nitrosalicylic acid**, 1.442, M-H_chebi, C7H5N1O5 |
| 182.0095 | 294.7596 | 5-nitrosalicylate, 1.0 |
| 194.0458 | 248.9994 | n-acetyl-5-aminosalicylic acid, 1.0, 6-hydroxy-3-succinoylpyridine, 1.0 |
| 204.1242 | 189.3015 | pantothenol, 1.0 |
| 215.0330 | 34.8192 | 2-c-methyl-d-erythritol 4-phosphate, 1.0 |
| 217.0300 | 34.7347 | diisopropyl sulfate, 1.0 |
| 224.8664 | 35.3805 | tetrathionate, 1.0 |
| 228.1607 | 442.6324 | n-decanoylglycine, 1.0 |
| 237.9310 | 209.2619 | 237.93104 mz/209.2619 s |
| 251.0962 | 390.4150 | girgensonine, 1.0 |
| 256.1922 | 491.1016 | n-lauroylglycine, 1.0 |
| 273.0078 | 297.9281 | ferulic acid 4-sulfate, 1.0, Isoferulic acid 3-sulfate, 1.0 |
| 273.1460 | 239.9128 | 273.14599 mz/239.91284 s** |
| 279.1638 | 563.6624 | 1-(3-chlorophenyl)-4-hexylpiperazine**, 0.128, M-H_chebi, C16H25Cl1N2 |
| 295.1588 | 551.8926 | ropinirole, 1.0 |
| 320.1906 | 475.7729 | (5Z,8Z,11Z,14Z,17Z)-icosapentaenoyl-containing glycerolipid**, 2.672, Cl_chebi, C20H291 |
| 328.2134 | 473.3893 | 6-keto-decanoylcarnitine, 1.0 |
| 331.2243 | 330.1762 | 331.22431 mz/330.17615s |
| 335.2233 | 425.4435 | 5s/15s/12r/11r/9s/8s/5/15-hpete, 1.0, lmfa03060044', 1.0, lmfa03060045', 1.0 |
| 347.2000 | 492.2968 | 7s,8s-dihode, 1.0, 5s,8r-dihode, 1.0, 15,16-dihode, 1.0, (±)-(e)-13-hydroxy-10-oxo-11-octadecenoic acid, 1.0 |
| 359.2557 | 381.5431 | 359.2557 mz / 381.5431 s ** |
| 405.1442 | 236.1413 | 2-[[2-[2-(2,3-dihydroindol-1-yl)-2-oxoethyl]-1-oxo-5-isoquinolinyl]oxy]acetic acid ethyl ester**, 3.826, M-H_chebi, C23H22N2O5 |
| 489.2031 | 226.6762 | 489.20314 mz /226.6762 s** |

**Table S6. Features important from both the RF and LR models with model usage.** Shown are the stability type, average RF feature important and sum of number of models the feature was used in across all three LR models over the 50 repeats.

| mz | rt | Stability type Building 1 | Stability type Building 2 | Stability type Building 3 | RF_feat_avg | RF_feat_std | LR_model_1_sum | LR_model_2_sum | LR_model_3_sum |
|---|---|---|---|---|---|---|---|---|---|
| 124.9913 | 34.4796 | stable | stable | stable | 0.0056 | 0.0018 | 40 | 0 | 50 |
| 140.9863 | 36.6597 | class 3 | stable | stable | 0.0113 | 0.0028 | 50 | 9 | 18 |
| 168.0302 | 183.5186 | stable | stable | stable | 0.0074 | 0.0026 | 50 | 50 | 2 |
| 182.0095 | 323.1458 | class 3 | class 3 | class 3 | 0.0056 | 0.0020 | 50 | 48 | 2 |
| 182.0095 | 294.7596 | class 3 | class 3 | stable | 0.0055 | 0.0020 | 8 | 48 | 1 |
| 194.0458 | 248.9994 | stable | stable | stable | 0.0145 | 0.0035 | 41 | 50 | 50 |
| 204.1242 | 189.3015 | class 3 | class 3 | stable | 0.0098 | 0.0028 | 48 | 29 | 1 |
| 215.0330 | 34.8192 | stable | stable | stable | 0.0079 | 0.0028 | 0 | 50 | 50 |
| 217.0300 | 34.7347 | stable | class 3 | stable | 0.0070 | 0.0028 | 0 | 49 | 22 |
| 224.8664 | 35.3805 | class 3 | class 3 | class 3 | 0.0048 | 0.0020 | 0 | 17 | 43 |
| 228.1607 | 442.6324 | stable | class 3 | class 3 | 0.0090 | 0.0029 | 50 | 45 | 3 |
| 237.9310 | 209.2619 | class 1 | class 3 | stable | 0.0085 | 0.0025 | 50 | 40 | 1 |
| 251.0962 | 390.4150 | class 3 | stable | stable | 0.0050 | 0.0019 | 50 | 45 | 2 |
| 256.1922 | 491.1016 | class 1 | class 3 | stable | 0.0145 | 0.0032 | 50 | 49 | 9 |
| 273.0078 | 297.9281 | stable | stable | class 3 | 0.0105 | 0.0028 | 50 | 41 | 2 |
| 273.1460 | 239.9128 | stable | stable | stable | 0.0087 | 0.0027 | 40 | 14 | 1 |
| 279.1638 | 563.6624 | class 3 | stable | stable | 0.0092 | 0.0026 | 50 | 13 | 2 |
| 295.1588 | 551.8926 | class 3 | stable | stable | 0.0051 | 0.0020 | 49 | 6 | 7 |
| 320.1906 | 475.7729 | class 1 | class 3 | stable | 0.0110 | 0.0030 | 50 | 9 | 50 |
| 328.2134 | 473.3893 | class 1 | class 3 | stable | 0.0087 | 0.0027 | 50 | 15 | 28 |
| 331.2243 | 330.1762 | stable | stable | stable | 0.0124 | 0.0030 | 50 | 27 | 1 |
| 335.2233 | 425.4435 | class 3 | stable | class 3 | 0.0077 | 0.0021 | 0 | 50 | 50 |
| 347.2000 | 492.2968 | class 3 | stable | stable | 0.0089 | 0.0032 | 50 | 50 | 1 |
| 359.2557 | 381.5431 | class 3 | stable | stable | 0.0075 | 0.0025 | 46 | 10 | 1 |
| 405.1442 | 236.1413 | stable | stable | stable | 0.0065 | 0.0022 | 50 | 42 | 1 |
| 489.2031 | 226.6762 | class 3 | class 3 | stable | 0.0081 | 0.0028 | 0 | 34 | 50 |

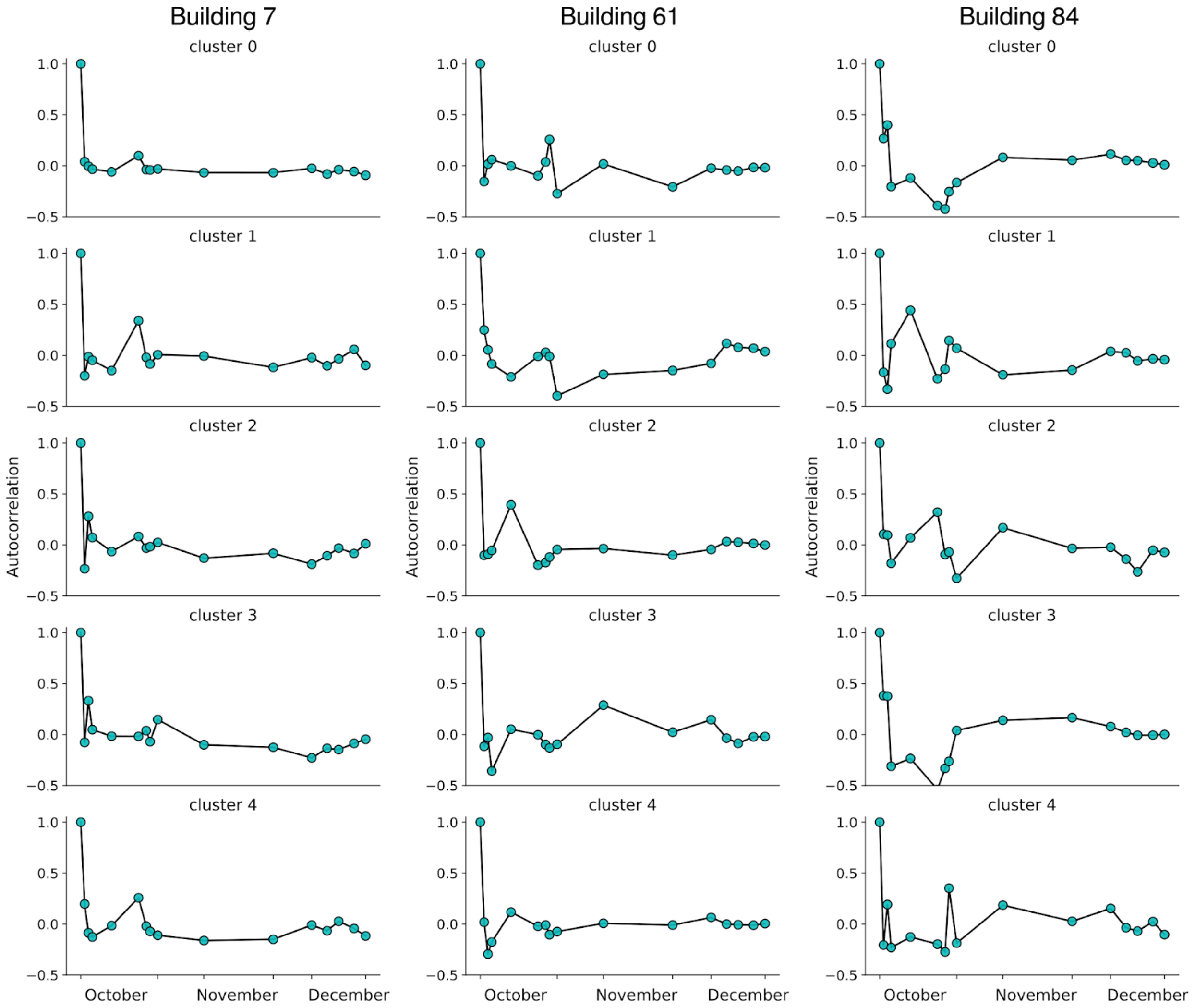


**Figure S8. Autocorrelation analysis of the 5 largest clusters for each of the 3 buildings showing rapid loss of autocorrelation.** Autocorrelation was calculated at all lag times for the data set and plotted on the x-axis where the showing where temporally the lag time would fall, however the values of lag time correspond to [0 … length of the time series] and not the actual dates. Autocorrelation was calculated with from statsmodels.tsa.stattools.acf
